# Highly sensitive genetically-encoded sensors for population and subcellular imaging of cAMP *in vivo*

**DOI:** 10.1101/2021.08.27.457999

**Authors:** Crystian I Massengill, Landon Bayless-Edwards, Cesar C Ceballos, Elizabeth R Cebul, Maozhen Qin, Matthew R Whorton, Bing Ye, Tianyi Mao, Haining Zhong

## Abstract

Cyclic adenosine monophosphate (cAMP) integrates information from diverse G protein-coupled receptors, such as neuromodulator receptors, to regulate pivotal biological processes in a cellular- and subcellular-specific manner. However, *in vivo* cellular-resolution imaging of cAMP dynamics in neurons has not been demonstrated. Here, we screen existing genetically-encoded cAMP sensors, and further develop the best performer to derive three improved variants, called cAMPFIREs. These sensors exhibit up to ten-fold increased sensitivity to cAMP and a corrected, cytosolic distribution. cAMPFIREs are compatible with both ratiometric and fluorescence lifetime imaging, and can detect cAMP dynamics elicited by norepinephrine at physiologically-relevant, nanomolar concentrations. Imaging of cAMPFIREs in awake mice reveals tonic levels of cAMP in cortical neurons that are associated with wakefulness, and are differentially regulated in different subcellular compartments. Furthermore, enforced locomotion elicits neuron-specific, bidirectional cAMP dynamics, in part, mediated by norepinephrine. Finally, cAMPFIREs also function in *Drosophila*, suggesting that they have broad applicability for studying intracellular signaling *in vivo*.

## Introduction

Cyclic adenosine monophosphate (cAMP) is a ubiquitous second messenger found in nearly all organisms, from bacteria to humans^1,2^. Three out of four major classes of G protein-coupled receptors (GPCRs) – those coupling to Gs, Gi and Gq proteins – function through the cAMP pathway^3–5^. In the mammalian brain, all major neuromodulators, such as norepinephrine, dopamine and acetylcholine, converge onto cAMP to regulate diverse brain function^3,5^. The dynamics of cAMP are both receptor-specific and cell-specific: the same extracellular ligand may elevate or decrease intracellular cAMP concentrations in different cells, depending on the cell-specific differential expression of receptor subtypes. Furthermore, cAMP dynamics within a cell may be spatiotemporally compartmentalized^6,7^.

Towards dissecting the spatiotemporal regulation of cAMP-dependent processes, over 50 genetically-encoded cAMP sensors and their improved variants have been developed^8,9^. These sensors have been used to report cAMP activity in a variety of *in vitro* preparations. However, the application of these sensors *in vivo*, particularly in neurons during animal behavior, remains difficult because the challenging *in vivo* imaging environment imposes high demands for sensor performance. No *in vivo* imaging of cAMP in individual neurons or in neuronal compartments have previously been achieved. As a result, the precise spatiotemporal dynamics of cAMP remain poorly understood in behaving animals.

Herein, we compared existing cAMP sensors^9–17^, both Förster resonance energy transfer (FRET)-based and intensity-based, under identical conditions in human embryonic kidney (HEK) 293 cells. The maximal dynamic ranges and the responses to a major neuromodulator, norepinephrine (NE) were determined. We identified the best performer, and further used structure-guided mutagenesis to engineer three enhanced variants, which we named <u>cAMP</u> <u>F</u>luorescence Imaging <u>R</u>eporters based on <u>E</u>pac (cAMPFIREs). These sensors bear improvements in two aspects. First, cAMPFIREs exhibit up to 10-fold increase in sensitivity to cAMP, enabling detection of cAMP dynamics in neurons elicited by physiologically-relevant, nanomolar concentrations of norepinephrine. Second, these sensors exhibit an even subcellular distribution, corrected from the uneven subcellular distribution and protein aggregation of the parental sensor. cAMPFIREs are compatible with both primary modalities of FRET imaging: ratiometric imaging and fluorescence lifetime imaging microscopy (FLIM). Importantly, these sensors allow for longitudinal imaging of bi-directional cAMP dynamics in mouse layer 2/3 (L23) cortical neurons *in vivo* with single-cell and subcellular resolutions. Using cAMPFIREs in the mouse somatosensory cortex, we found a tonic level of cAMP that was dependent on wakefulness, and was in part mediated by adrenergic receptors in a neuronal compartment-specific manner. At the population level, animal locomotion triggered cell-specific, up or down regulations of cAMP in different L23 neurons, which were mediated in part by different adrenergic receptors. *In situ* live imaging of cAMP dynamics in neurons of *Drosophila* larva was also demonstrated. cAMPFIREs may enable *in vivo* interrogation of cAMP dynamics underlying animal behavior, and have broad application in other paradigms where cAMP is involved.

## Results

### Screening existing cAMP sensors under identical conditions

Learning from the development of genetically-encoded calcium and protein kinase A (PKA) sensors^18–20^, we first screened for the best existing cAMP sensors under identical conditions in HEK-293 cells. Eight sensors that were recently developed or frequently used were tested (Figs. 1a and 1b)^9–17^, including both FRET-based and intensity-based sensors. Each sensor was assessed for both its response to a physiological ligand, norepinephrine (1 μM), which stimulates cAMP production through the β-adrenergic receptor pathway, and its maximal dynamic range, as elicited by bath application of the adenylyl cyclase agonist forskolin (25 μM) and phosphodiesterase inhibitor IBMX (50 μM). To compare across imaging modalities, the response amplitudes were normalized to their respective baselines (i.e., ΔRatio/Ratio_0_ or ΔF/F_0_) (Figs. 1c – 1e).

**Fig. 1.**
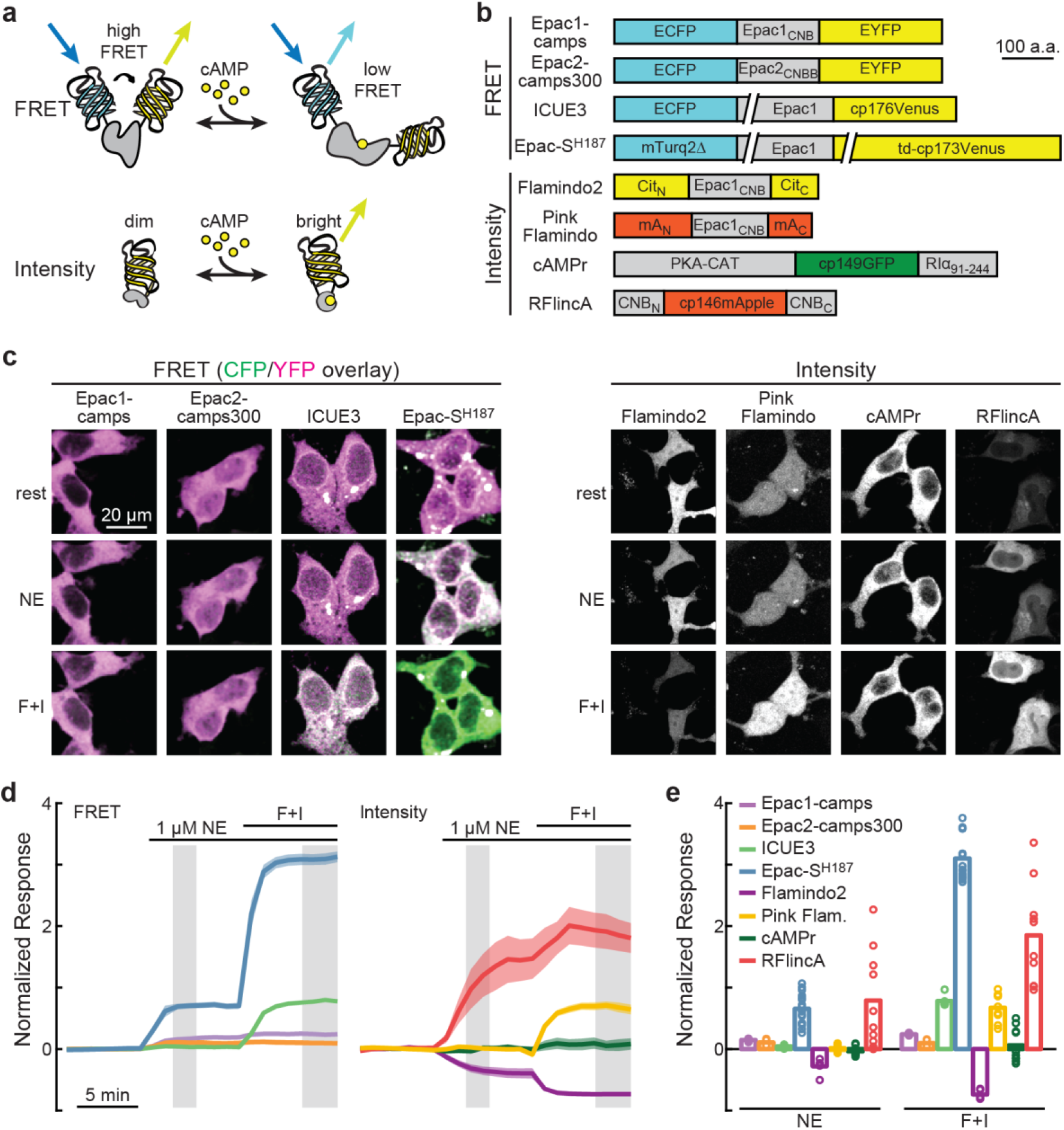
Screening of existing cAMP sensors. **a**, General schematic of FRET-based and intensity-based cAMP sensors. **b**, Schematics of the cAMP sensors tested. Epac2_CNBB_: CNB-B domain of Epac2; Cit_N_ and Cit_C_: residue 1–144 and 146–238 of Citrine, respectively; mA_N_ and mA_C_: residues 1–150 and 151–246 of mApple, respectively; PKA-CAT: PKA catalytic subunit α; RIα_91-244_: residue 91–244 of PKA regulatory subunit Iα; CNB_N_ and CNB_C_: N-terminal and C-terminal portions of the CNB domain of PKA regulatory subunit Iα, respectively; cp: circularly permutated. **c**, Representative images of the tested sensors at rest, after stimulation with NE (1 μM), and followed by 25 µM forskolin and 50 µM IBMX (F + I). CFP and YFP ratiometric images are shown as a pseudocolored green (CFP) and magenta (YFP) overlay. To compare across cells, for ratiometric images, the same ratio of intensity setting was used for image display across sensors; whereas intensity images were adjusted according to the brightest state. **d** & **e**, Averaged response (ΔR/R_0_, or ΔF/F_0_) traces (**d)** and quantifications at the grayed time window in panel **d** (**e)** of each sensor. From left to right, n (cells) = 5, 4, 12, 16, 5, 10, 10, 10. Dark line and shaded area indicate average and s.e.m. respectively. Bars indicate mean.

Among all sensors tested, Epac-S^H187^ (Fig. 1b; ref 14) was selected for further development for the following reasons: First, it exhibited the largest dynamic range and second-largest response to norepinephrine (ΔRatio/Ratio_0_ = 0.68 ± 0.05 for norepinephrine, and 3.11 ± 0.09 for forskolin/IBMX) (Figs. 1d and 1e). Second, compared to other alternatives, Epac-S^H187^ exhibited fast kinetics and consistent responses across experiments (Figs. 1d and 1e). Third, Epac-S^H187^ uses mTurquoise2 as the donor fluorophore, which exhibits single exponential fluorescence lifetime decay. This makes the sensor also suitable for FLIM imaging^9,21,22^, which has advantages when imaging slow (seconds to minutes) signaling events *in vivo*, such as in the brain during behavior. However, Epac-S^H187^ also exhibited two limitations. First, its response to norepinephrine was only 22% of its maximal dynamic range, suggesting room for improvement in sensor sensitivity (Figs. 1d and 1e). Increasing cAMP sensitivity would maximize the signal amplitude and therefore the signal-to-noise ratio under physiological, *in vivo* imaging conditions. Second, the sensor adopted an uneven, perinuclear distribution in cells^23,24^. Correcting this uneven distribution would enhance sensor use at desired locations and reduce potential impacts on cellular function.

### Engineering the sensor to correct for uneven subcellular distribution

We first dealt with the uneven distribution of Epac-S^H187^ in cells. An ideal sensor should distribute evenly in cells unless specifically targeted. However, the parental protein of this sensor, Epac1, is known to adopt specific subcellular localizations on the membrane and at the perinuclear structure via its N-terminal Dishevelled, Egl-10, and Pleckstrin (DEP) domain and C-terminal nuclear pore localization (NPL) sequence, respectively (Fig. 2a)^23–25^. In Epac-S^H187^, the DEP domain has been removed^26^; however, the NPL domain remains (Figs. 2a and 2b) and preferentially localizes the sensor to perinuclear locations (Figs. 2c and 2d, and Extended Fig. S1). This subcellular distribution is not only undesirable from an experimental perspective, but may also negatively impact cellular function: the sensor no longer contains a functional Epac protein^26^, and occupying the location of endogenous Epac could potentially cause dominant negative effects.

**Fig. 2.**
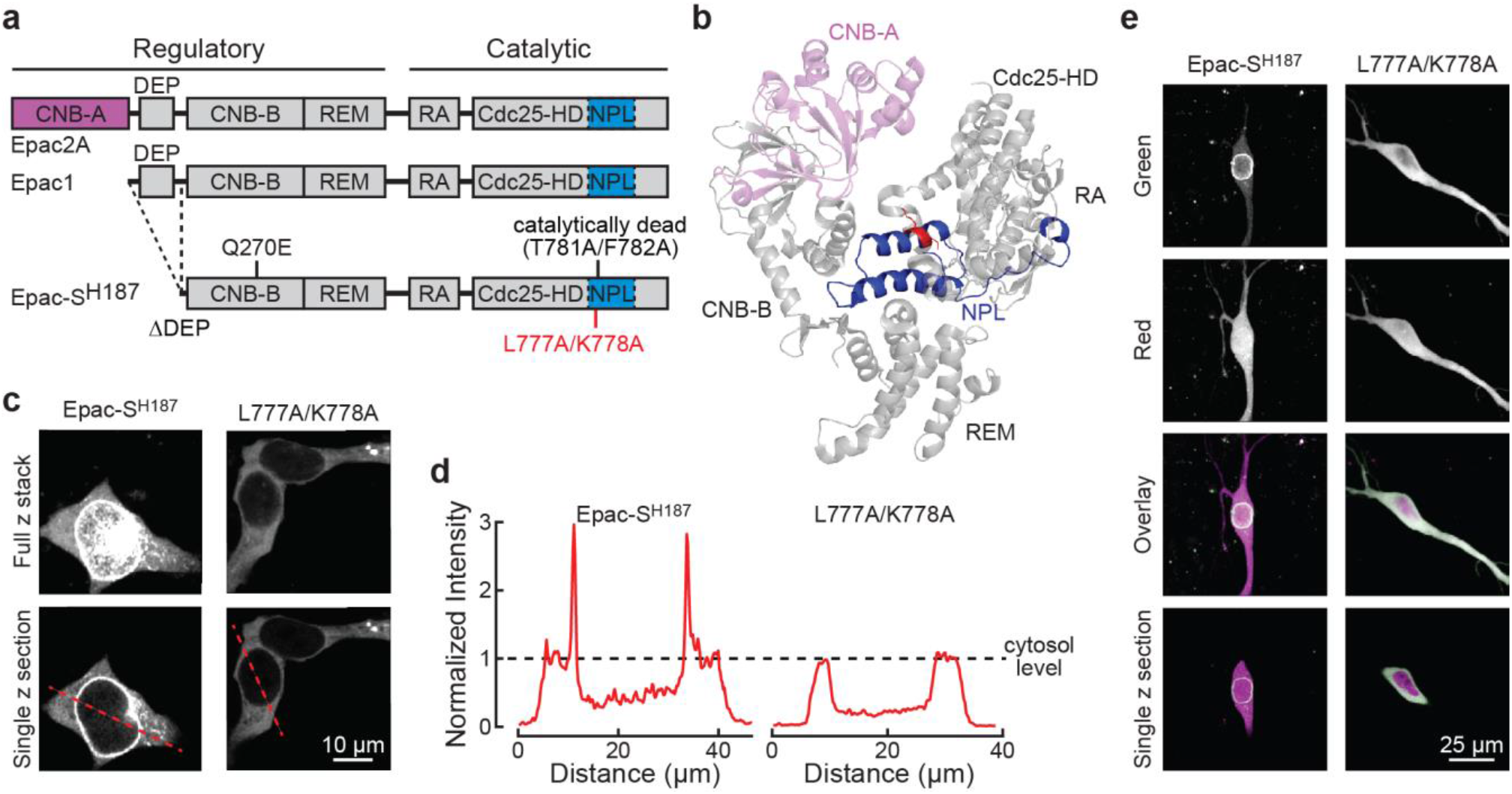
Correcting the perinuclear localization of Epac-S^H187^. **a,** Schematic domain organization of Epac2A, Epac1, and Epac-S^H187^. CNB, cyclic nucleotide binding domain; DEP, Dishevelled, Egl-10, Pleckstrin and domain; REM, Ras-exchange motif; RA, ras-association domain; CDC25-HD, CDC25 homology domain; NPL, nuclear pore localization sequence. **b,** The crystal structure of Epac2A in the cAMP-unbound conformation (Protein data bank #2byv). **c,** Representative two-photon images of Epac-S^H187^ and the L777A/K778A mutant expressed in HEK cells. **d,** Quantification of fluorescence intensity along the indicated dashed line in panel **c**, normalized to cytosolic average. **e,** Representative two-photon images of nuclear pore localization of Epac-S^H187^ and the cytosolic distribution of the L777A/K778A mutant in cultured CA1 neurons.

The NPL sequence has been previously narrowed to a 74-residue sequence at the C-terminus of Epac1 (blue portion in Fig. 2b)^24^. It was also reported that, although the homologous Epac2 protein also contained a functional NPL sequence, it was sterically blocked by the CNB-A domain (light magenta in Fig. 2b)^24,27^. We hypothesized that exposed residues within the NPL sequence spatially facing the CNB-A domain may mediate the perinuclear localization of Epac. An analysis of the crystal structure of Epac2^27^ identified 12 candidate residues that were also conserved in Epac1 (Extended Data Fig. 1a). By systematic mutagenesis of these residues to alanine, two mutants, F772A/L775A and L777A/K778A, were found to exhibit disrupted perinuclear localization (Figs. 2c and 2d, and Extended Data Fig. 1b). Among the two, the F772A/L775A mutant largely lost its responsiveness, but the L777A/K778A mutant exhibited increased response amplitudes compared to the parental sensor (41% increase to 1 µM NE; Extended Data Fig. 1c). This mutant also exhibited an even, cytosolic distribution when expressed in CA1 neurons in cultured hippocampal slices (Fig. 2e) and in L23 pyramidal neurons in acute brain slices (Extended Data Fig. 1d).

In addition, we found that Epac-S^H187^ exhibited a high tendency to form aggregates. Consistent with a previous report, this tendency was already lowered compared to a variant that used only a single cpVenus as the acceptor (Extended Data Fig. 2)^28^. The L777A/K778A mutation further reduced the number of aggregates by ∼5 fold (Extended Data Fig. 2), suggesting that the aggregation is related to the perinuclear localization of the sensor.

### Increasing the response sensitivity of the sensor

As mentioned earlier, the response of Epac-S^H187^ to norepinephrine was 22% of its dynamic range (Fig. 1d and 1e), suggesting an insufficient sensitivity to physiological concentrations of cAMP. Indeed, recombinant Epac-S^H187^ were found to exhibit a cAMP EC50 of ∼3.8 μM^14^, which is several folds higher than predicted cAMP concentrations in cells (∼1 μM) and the cAMP EC50 of PKA^29–31^. We used a two-pronged approach to increase the sensitivity of Epac-S^H187^. First, we examined the crystal structures of the homologous Epac2 protein^32^, focusing on the CNB-B domain, which is equivalent to the only CNB domain in Epac1 (Fig. 2a and Extended Data Fig. 3a). Ten candidate residues at or near the cAMP binding site were identified that might affect the affinity (Extended Data Fig. 3a). The first round of mutagenesis found that one candidate mutation (M312F) nearly completely abolished the sensor response, indicating that it was at a position critical to cAMP binding. Further random mutagenesis at this position identified several mutants with increased norepinephrine response, with M312L providing a response 265 ± 15% as large as the original sensor (p<0.001; asterisk in Fig. 3b). Second, we used a candidate approach and tested mutations known to enhance ligand affinities in other CNB domain-containing proteins, specifically the cyclic nucleotide-gated (CNG) channels. Three candidate residues were identified and were aligned to F268, D306, and E325 in the Epac1 CNB domain (Extended Data Fig. S3b)^33,34^. Mutagenesis revealed that several mutations at the E325 position increased the response to norepinephrine. The best performer, E325T, exhibited a response 367 ± 13% as large as that of the parental sensor (triangle in Fig. 3b).

**Fig. 3.**
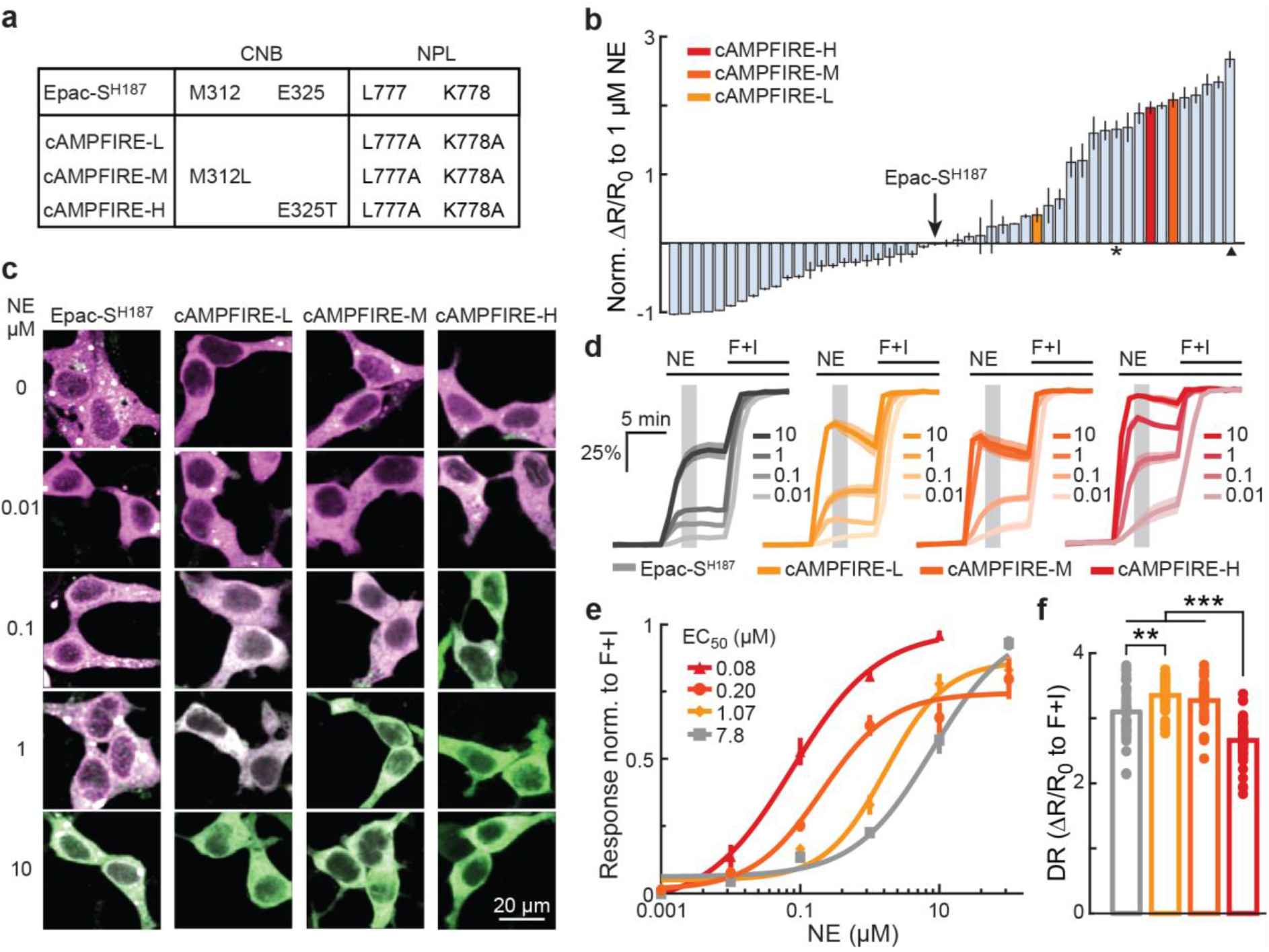
Increasing the cAMP sensitivity of Epac-S^H187^ to derive cAMPFIREs. **a**, Table illustrating the mutations in each cAMPFIRE variant as compared to the parental sensor. **b**, The fold changes in the response of screened variants to 1 μM norepinephrine relative to Epac-S^H187^. n = 2 – 20 cells each. Bars and error bars indicate mean and s.e.m, respectively. **c – e**, Representative ratiometric images (**c**), response traces normalized to the corresponding responses elicited by forskolin and IBMX (**d**), and dose-response curves to norepinephrine fitted by the Hill equation (**e**), of Epac-S^H187^ and cAMPFIRE-L, M, and H in HEK cells. n = 6 – 20 for each sensor and concentration data point. In panel **d**, dark lines and shaded area, and in panel **e**, points and error bars indicate mean and s.e.m respectively. **f**, Response amplitudes of each sensor elicited by forskolin and IBMX. DR: dynamic range. From left to right, n (cells) = 60, 53, 49, and 49. Bars represent mean. ***: p < 0.001 and **: p < 0.01, dF = 3, one-way ANOVA, Tukey-Kramer post-hoc test.

We combined the “delocalization” mutations (L777A/K778A) either without or individually with M312L or E325T to derived three improved sensors (Fig. 3a). Because Epac-S^H187^ was named using a construct number internal to the producer’s lab, we renamed our variants cAMPFIRE-L, -M, and -H, standing for low, medium and high affinity, respectively (Figs. 3a and 3b). To better quantify the degree of increased sensitivity to norepinephrine, we determined the norepinephrine dose-response curve of cAMPFIREs and Epac-S^H187^. To control for potential adaptation of the cAMP pathway to multiple stimulations, only one dose of norepinephrine was given in each experiment, and responses across experiments were normalized to the corresponding maximal response elicited by forskolin and IBMX before being pooled together (Figs. 3c and 3d). Norepinephrine has been shown to trigger physiological responses at a concentration of ∼ 0.02 µM^20^. However, Epac-S^H187^ exhibited an EC50 of 7.8 ± 3.0 µM (fitted value ± 95% confidence interval; Fig. 3c – 3e), over two orders of magnitude higher. cAMPFIRE-L, -M, and -H showed 7, 38, and 101-fold increases, respectively, in sensitivity to norepinephrine (Fig. 3e). As little as 10 nM norepinephrine could be detected. The dynamic ranges of cAMPFIREs, as elicited by forskolin/IBMX, only changed moderately compared to Epac-S^H187^, with cAMPFIRE-L exhibiting a ∼8% increase and cAMPFIRE-H exhibiting a ∼14% decrease (ΔRatio/Ratio_0_ = 3.33 ± 0.04, 3.27 ± 0.05, and 2.64 ± 0.05 for cAMPFIRE-L, -M, and -H, respectively, cf. 3.11 ± 0.09 for Epac-S^H187^; Fig. 3f). The moderately decreased dynamic range in cAMPFIRE-H was likely because its much increased sensitivity resulted in sensor activation at basal cAMP levels, as reflected by its elevated baseline (Extended Data Fig. 4). Similar increases in sensitivity were observed when the responses were measured using two-photon FLIM (2pFLIM; Extended Data Fig. 5), which we used for tissue and *in vivo* imaging.

### Characterizing the specificity and properties of cAMPFIREs

With their improved sensitivity and corrected subcellular distribution, cAMPFIREs hold the promise of becoming the next-generation cAMP sensors. We therefore systematically characterized their properties. The response of cAMPFIRE-L to norepinephrine was abolished when a key residue essential for cAMP binding was mutated (Fig. 4a)^35^, suggesting that the detected response was specifically due to cAMP binding. Moreover, the response to norepinephrine was reversible (Fig. 4b), and it was completely blocked by the β-adrenergic receptor antagonist propranolol (1 µM) (Fig. 4b).

**Fig. 4.**
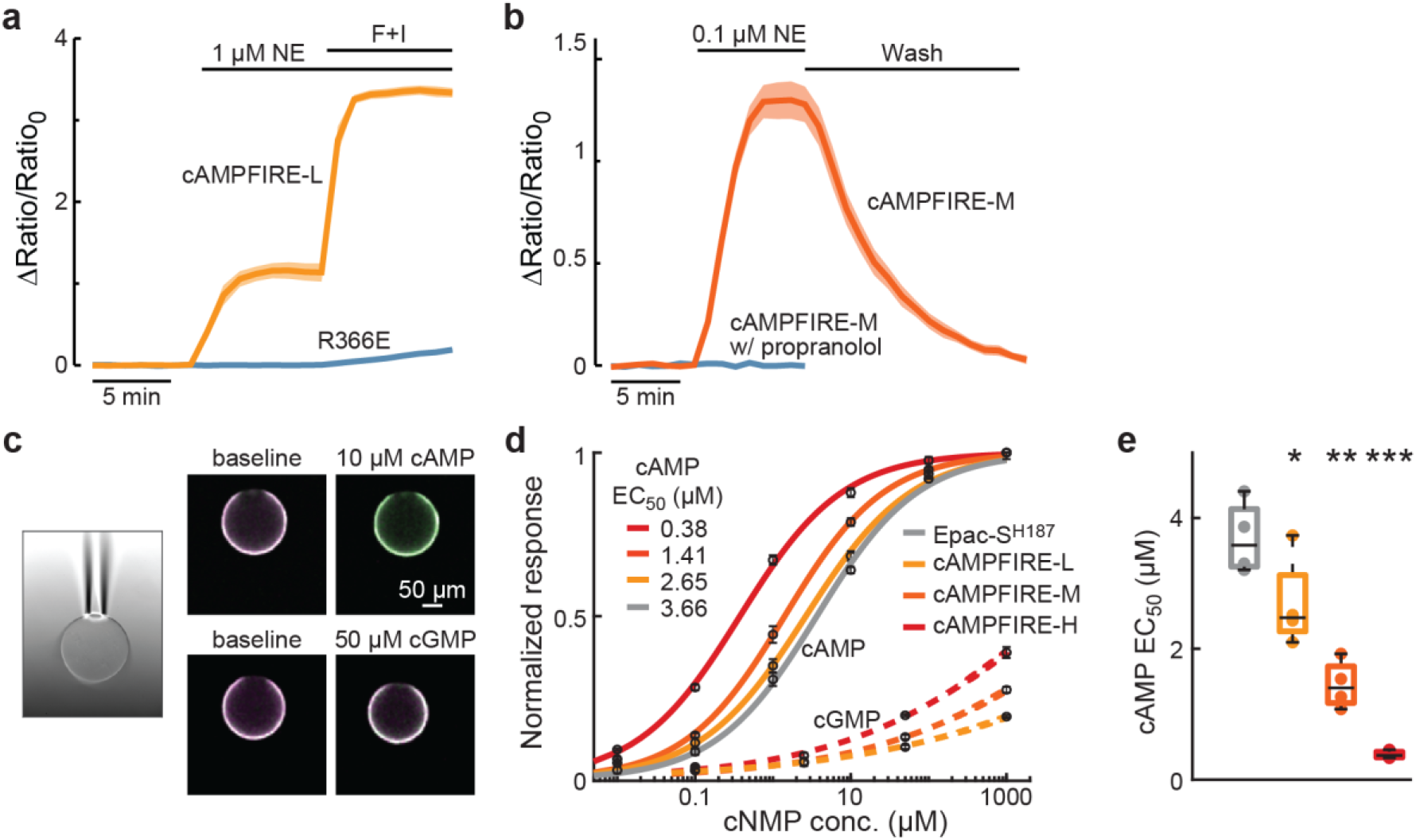
Characterization of cAMPFIREs expressed in HEK cells. **a**, Response traces of cAMPFIRE-L and its cAMP-binding mutant (R366E) to the indicated stimulus. n (cells) = 8 and 11 for cAMPFIRE-L and R366E, respectively. **b**, Response traces of cAMPFIRE-M to 0.1 µM norepinephrine followed by wash, in the absence or presence of propranolol. n (cells) = 8 for both. Dark lines and shaded area indicate mean and s.e.m., respectively. **c** – **e**, Representative images (**c**), dose-response curves to cAMP and cGMP (**d**), and cAMP EC50s (**e**) of the cAMPFIRE sensors compared to the parental sensor. Asterisks indicate the significance levels compared to Epac-S^H187^. n = 4 beads each. In panel **d**, points and error bars indicated mean and s.e.m., respectively. In panel **e**, boxes indicate 25th and 75th percentile, with black lines indicating median and whiskers indicating total range of data, not including outliers. *: p < 0.05, **: p < 0.01, and ***: p < 0.001, dF = 6, two-tailed unpaired Student’s t-test on log-transformed data.

While sensor responses measured up to this point had used norepinephrine as the physiological stimulus, it is also critical to directly measure the cAMP affinity and specificity. To this end, we tagged each sensor with a myc tag at its C terminus and immunoprecipitated the sensor to agarose beads using an anti-myc antibody (see **Methods**). The beads were then fixed in the perfusion chamber for imaging using a suction pipet (Fig. 4c), allowing the determination of the dose-response relationship to cAMP, and, importantly, to the structurally similar analog cGMP, which is also a major intracellular second messenger (Fig. 4d). The parental sensor exhibited a cAMP affinity of 3.7 ± 0.3 µM, consistent with the literature^14^. cAMPFIRE-L, -M, and –H all showed significantly increased sensitivity to cAMP, with cAMPFIRE-H exhibiting a ∼10-fold increase (EC50 = 2.7 ± 0.4, 1.45 ± 0.18, and 0.38 ± 0.03 for cAMPFIRE-L, -M, -H, respectively; Figs. 4d and 4e). In contrast, cAMPFIREs exhibited little response to cGMP. At the highest concentration we tested (1000 µM), which is >100 fold higher than physiological concentrations of cGMP, all sensors responded well below 50% of their respective maximal responses. By fitting the partial curve and forcing the maximal normalized response to be 1, the EC50s for cGMP was estimated to be 56,000 ± 16,000, 30,100 ± 19,000, and 5,000 ± 550 µM for cAMPFIRE-L, M, and H, respectively, all over four orders of magnitude higher than the corresponding affinity for cAMP. Taken together, cAMPFIREs specifically detect cAMP with increased sensitivity up to the submicromolar range.

### cAMPFIREs detect neuromodulatory cAMP dynamics in brain tissue without affecting neuronal function

With the goal to image these sensors *in vivo* in the brain, we next asked whether cAMPFIREs can detect cAMP increases elicited by norepinephrine in neurons within brain tissue. We imaged cAMPFIREs in the apical dendrites of CA1 neurons in cultured hippocampal slices, as transfected using the biolistic method, and in L23 pyramidal neurons in acute brain slices of mouse somatosensory cortices, as transfected using *in utero* electroporation (IUE; Extended Data Fig. 6). For these experiments, we used 2pFLIM to measure FRET, which offers advantages for quantifying FRET in brain tissue and *in vivo*, including being quantitative, resistant to wavelength-dependent light scattering, and less sensitive to parameter changes in imaging conditions^9,21,22^. cAMPFIREs exhibited substantially increased response amplitudes compared to Epac-S^H187^ in CA1 neurons of organotypic cultured hippocampal slices, especially when norepinephrine was used (compared to Epac-S^H187^, the response were 195 ± 25%, 180 ± 19%, and 179 ± 15% for cAMPFIRE-L, M, and H, respectively; for all cAMPFIREs, p < 0.01, cf. Epac-S^H187^) (Figs. 5a and 5b). These responses were reversible and could be blocked by the bath application of the β antagonist propranolol (Extended Data Fig. 7a). Similarly, in L23 pyramidal neurons in acute brain slices, cAMPFIRE-L and -M responded robustly to norepinephrine at concentrations as low as 10 nM, which is known to be sufficient for physiological responses^20^ (Fig. 5c and 5d). These responses were ∼3-fold as large as those from the parental sensor under the same conditions (336 ± 71% and 290 ± 77% for cAMPFIRE-L and M, respectively). When comparing across cell types, the basal lifetime of cAMPFIRE-M increased from HEK cells to CA1 neurons and further to L23 neurons, resulting in reduced dynamic range (Extended Data Fig. 7b). This suggests that L23 neurons exhibit a cell type-specific tonic level of cAMP, which is sufficient to activate cAMPFIRE-M even at rest (see also ref 20, 31). Therefore, the most sensitive cAMPFIRE-H was not further tested in L23 neurons. These data indicate that the cAMPFIRE variants indeed enable more sensitive detection of cAMP in neurons.

**Fig. 5.**
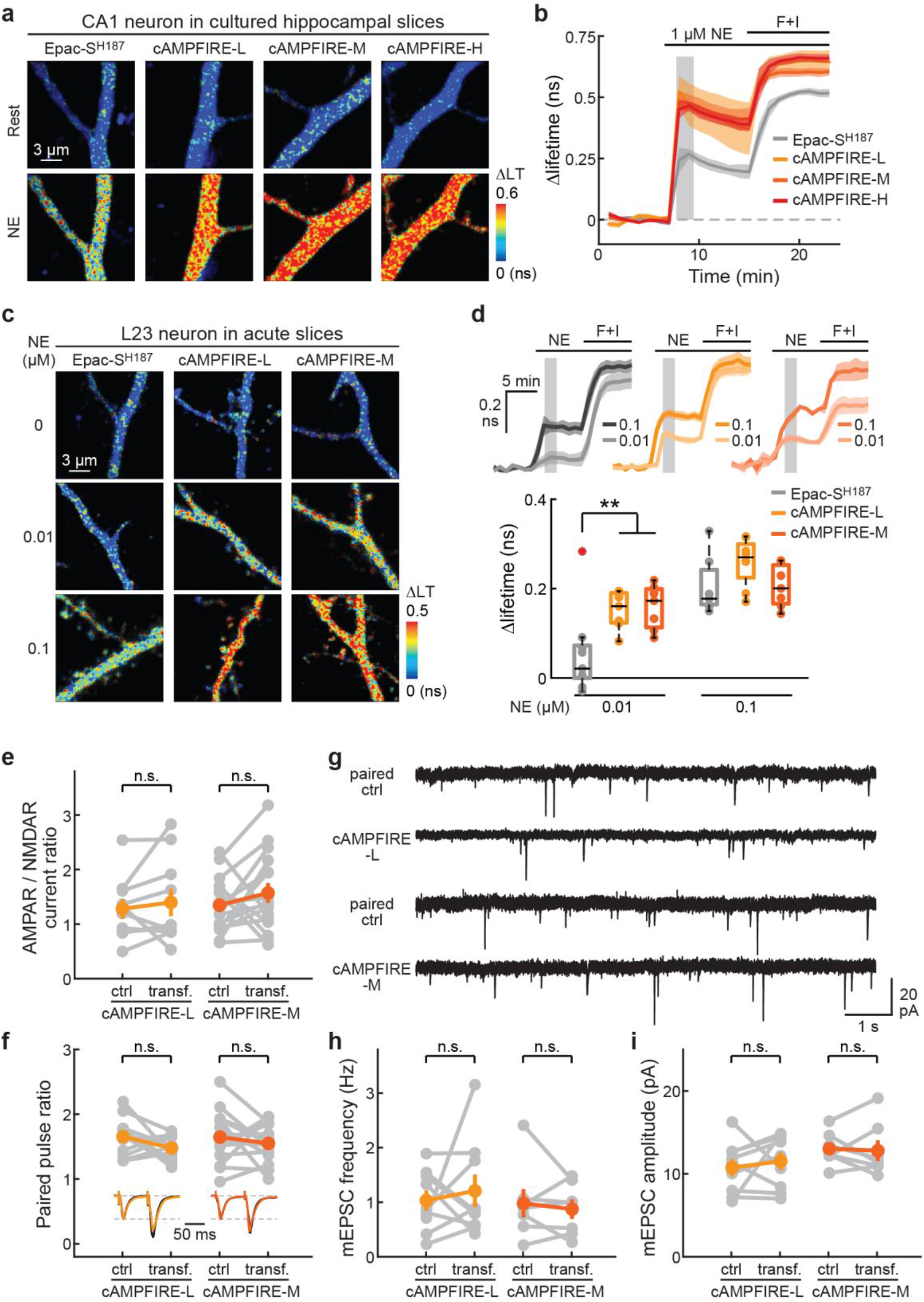
Characterization of cAMPFIREs in neurons from brain slices. **a** & **b**, Representative 2pFLIM images (**a**) and response traces (**b**) of cAMPFIREs and their parental sensor to the indicated stimuli. n = 5, 3, 8, and 5 for Epac-S^H187^, cAMPFIRE-L, -M, and -H, respectively. Dark lines and shaded area indicate mean and s.e.m, respectively. Compared to Epac-S^H187^, p < 0.01 for cAMPFIRE-L and -M and p < 0.001 for cAMPFIRE-H at shaded time points, two-tailed unpaired Student’s t-test, dF = 6, 11, 8 for cAMPFIRE-L, -M and -H, respectively. LT: lifetime. **c** & **d,** Representative 2pFLIM images (**c**), and response traces and quantifications (**d**) of the indicated sensors to the indicated stimulus. From left to right on the lower plot, n (neurons) = 9, 7, 7, 7, 8, and 7. **: p < 0.01, dF = 14 for both, two-tailed unpaired Student’s t-test. Dark lines and shaded area in the upper plot indicate mean and s.e.m., respectively. In lower plot, boxes indicate 25^th^ and 75^th^ percentile, with black lines indicating mean and whiskers indicating total range of data, excluding outliers. **e** – **g**, Quantification of the AMPAR/NMDAR current ratio (**e**; n = 10 for cAMPFIRE-L and 16 for cAMPFIRE-M), paired pulsed ratio (**f**; n = 11 for -L and 15 for -M), and the example traces (**g**), frequency (**h**), and amplitudes (**i**) of mEPSCs (n = 9 for -L and 7 for -M) of CA1 neurons transfected with cAMPFIRE-L and cAMPFIRE-M compared to adjacent untransfected control neurons. n.s.: not significant, paired two-tailed Student’s t-test. Colored points and error bars indicate mean and s.e.m., respectively.

To examine whether cAMPFIREs may potentially affect neuronal functions, we performed whole-cell recording of CA1 neurons in cultured hippocampal slices expressing cAMPFIRE-L or –M, and compared the neuronal properties to adjacent untransfected neurons (Figs. 5e – 5i, and Extended Data Fig. 8). The parameters tested included evoked AMPA and NMDA receptor currents (Extended Data Fig. 8), AMPAR/NMDAR current ratios (Fig. 5e), paired-pulse facilitation (Fig. 5f), the frequency and amplitude of miniature excitatory postsynaptic currents (Figs. 5g – 5i), as well as resting membrane potential and action potential frequencies (data not shown). The results showed that neither the transfection of cAMPFIRE-L nor -M affected any of these parameters. Thus, the expression of cAMPFIREs did not appear to alter neuronal function under our conditions. Overall, our results indicate that cAMPFIREs enhance detection of norepinephrine-elicited cAMP dynamics with high sensitivity that was not attainable using previous sensors.

### cAMPFIREs enable longitudinal and subcellular imaging of cAMP in awake mice

To test whether cAMPFIREs are sufficient for *in vivo* imaging of cAMP in awake mice, we used *in utero* electroporation to introduce cAMPFIRE-L or -M to L23 pyramidal neurons in the somatosensory cortex (Extended Data Fig. 6), and performed 2pFLIM through a craniotomy-implanted optical window in awake, head-fixed mice^36^. Individual neurons can be clearly resolved and be repeatedly imaged over months (Fig. 6a). The basal lifetimes of individual neurons were consistent across weeks (Fig. 6b). Interestingly, compared to acute slices, where neuromodulators have been washed away, both cAMPFIRE-L and cAMPFIRE-M exhibited higher basal lifetime *in vivo*, suggesting an additional tonic level of cAMP (Fig. 6c). Neuronal dendrites could also be resolved, and they exhibited higher lifetime compared to their corresponding somas (Fig. 6d), indicating differential tonic levels of cAMP between these neuronal compartments. These result indicates that cAMPFIREs are sufficient for subcellular-resolution imaging *in vivo*.

**Fig. 6.**
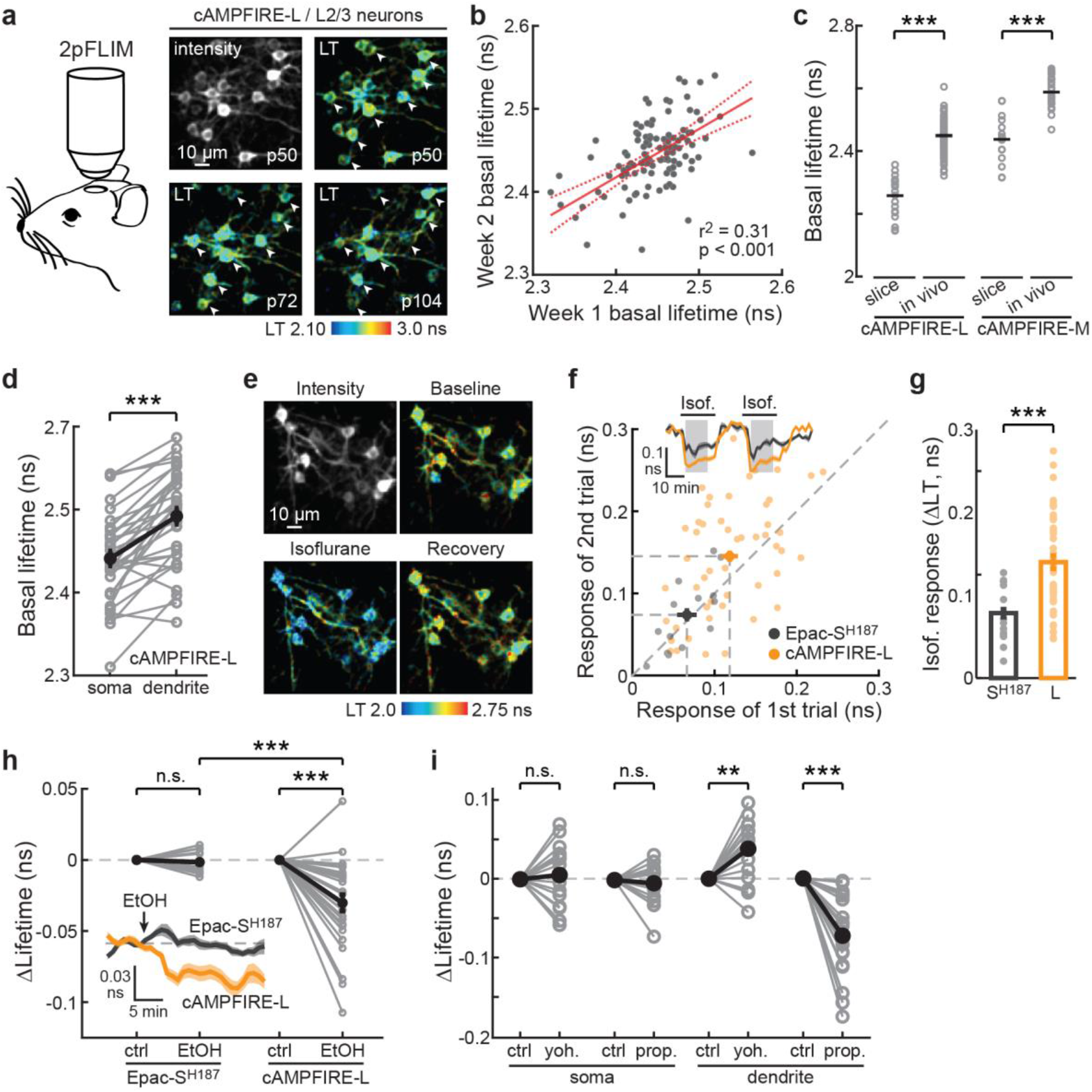
cAMPFIREs enables longitudinal *in vivo* imaging. **a**, Schematic and representative longitudinal images of 2pFLIM through a cranial window in awake mice. LT: lifetime. Arrowheads indicate the same cells across images. **b**, Correlation of basal lifetimes of the same cells imaged 5 – 8 days apart. n (cells) = 113. Solid line indicates fit and dashed lines indicate 95% confidence interval. **c**, Basal lifetimes of cAMPFIRE-L and M of individual neurons *in vivo* compared to those in acute slices. From left to right, n (cells) = 20, 151, 17, and 24. Horizontal black lines indicate mean. ***: p < 0.001, two-tailed Welch test, dF = 11.7 and 11.2 for cAMPFIRE-L and -M, respectively. **d**, Comparison of basal lifetimes between somas and dendrites. n (cells) = 35. Black points and error bars indicate mean and s.e.m. respectively. ***: p < 0.001, dF = 33, paired two-tailed Student’s t-test. **e**, Representative images of cAMPFIRE-L before (Baseline), during (Isoflurane), and after (Recovery) isoflurane administration. **f** & **g**, Average response traces (inset) and correlation (**f**), and collective response amplitudes of the first trial (**g**) of Epac-S^H187^ and cAMPFIRE-L in L23 neurons responding to two sequential isoflurane trials. The line of equality (not from fitting) is also shown. n (cells) = 14 for Epac-S^H187^, and 50 for cAMPFIRE-L. Bars and error bars indicate mean and s.e.m., respectively. ***: p < 0.001, dF = 72.3, two-tailed unpaired Welch test. **h**, Average response traces (inset) and collective response amplitudes of Epac-S^H187^ and cAMPFIRE-L in L23 neurons responding to EtOH injection. n (cells) = 12 for Epac-S^H187^, and 28 for cAMPFIRE-L. Black points and error bars in main figure and dark lines and shaded area in inset indicate mean and s.e.m., respectively. Within a sensor, ***: p < 0.001, unpaired, two-tailed Welch test. Across sensor, ***: P < 0.001, paired, two-tailed Student’s t-test. **i**, Soma and dendrite-specific cAMPFIRE-L response to yohimbine (yoh.) and propranolol (prop.). n (cells) = 16 for yohimbine, and 17 for propranolol. Black points indicate mean. **: p < 0.01, ***: p < 0.001, paired, two-tailed Student’s t-test.

It has been previously shown that animal wakefulness is associated with elevated levels of neuromodulators, such as norepinephrine, in the cortex^20,37^. We therefore asked whether the tonic cAMP in individual neurons was regulated by wakefulness using an established manipulation – anesthesia by isoflurane. We found that a moderate concentration of isoflurane (1.5%) significantly reduced cAMP levels as measured using cAMPFIRE-L in neuronal somas (Fig. 6e and 6f). This response to isoflurane was reversible and repeatable (Fig. 6f). Notably, the response amplitudes of cAMPFIRE-L were significantly larger than those detected by Epac-S^H187^ (88 ± 21% larger; Fig. 6g). We also tested ethanol, which overall depresses the brain function^38^. Intraperitoneal injection of ethanol (1.28 g/kg) resulted in decreased cAMP levels in cortical neurons as detected using cAMPFIRE-L (Fig. 6h). In contrast, Epac-S^H187^ failed to detect any differences under the same conditions, illustrating that the higher sensitivity of cAMPFIREs enables detection of previously undetectable cAMP dynamics.

To test whether the basal cAMP tone in L23 neurons is mediated by norepinephrine, we injected mice intraperitoneally with the α2 adrenergic antagonist yohimbine or the β antagonist propranolol in separate experiments. Interestingly, yohimbine injection resulted in increased cAMP concentrations in neuronal dendrites, but with little effect on neuronal somas (Fig. 6i). In contrast, propranolol injection decreased dendritic cAMP, and also with little effect on somatic cAMP (Fig. 6i). These results have several implications: First, cAMPFIREs are sufficient to detect bidirectional cAMP dynamics *in vivo*; second, neuronal dendrites and somas are unique cAMP compartments *in vivo* and may exhibit distinct cAMP dynamics; and third, a basal level of norepinephrine likely exists in the cortex of awake mice that results in a push-and-pull control of the basal cAMP activity in neuronal dendrites.

### cAMPFIREs reveal cell-specific cAMP dynamics during mouse locomotion

To evaluate whether cAMPFIREs may be used to detect cAMP dynamics elicited by animal behaviors, we used the mouse head-fixed locomotion paradigm (Fig. 7a)^36,39,40^. We imaged population responses of L23 neurons to enforced locomotion, and found that individual neurons, sometimes in the same field of view, exhibited distinct, cell-specific cAMP dynamics: some with positive cAMP responses, some with negative responses, and others with little response (Figs. 7b and 7c). The responses were reversible and repeatable for the same cell (Fig. 7c). As a result, the responses were much more correlated across trials within a cell than across cells (Fig. 7d, see **Methods**). Consistent with the empirical observation of this heterogeneity, three distinct clusters were revealed when averaged individual cell responses were analyzed using hierarchical cluster analyses (Fig. 7e and 7f).

**Fig. 7.**
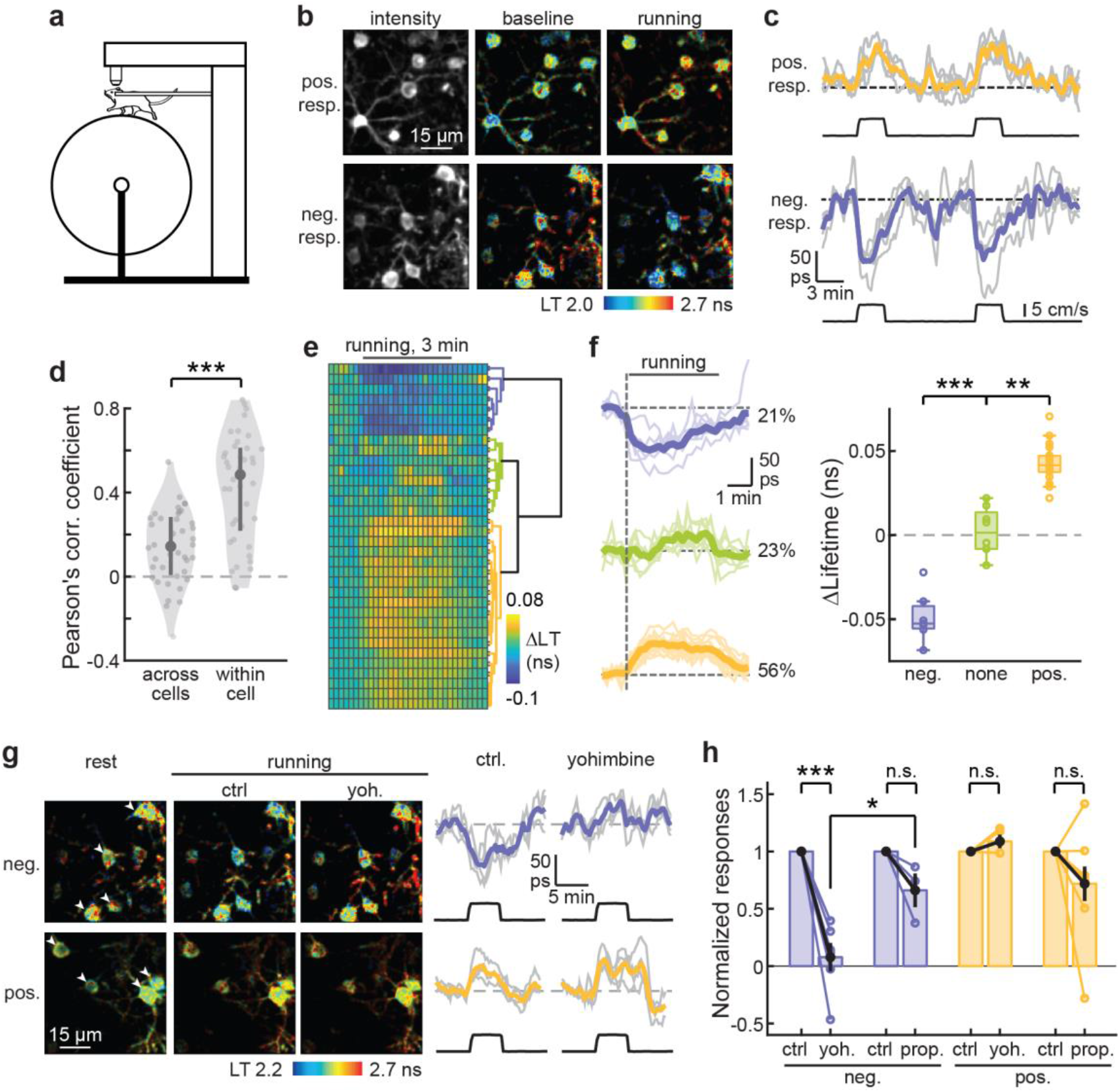
cAMPFIRE-L detects heterogeneous cAMP response to enforced running. **a**, Schematic of the enforced running behavioral paradigm. **b** & **c**, Representative images (**b**) and example response traces and their averages (**c**) of cAMPFIRE-L in positively and negatively responding layer 2/3 cells in response to enforced running. Colored lines indicate average response. Black lines indicate running speed. **d**, Quantification of between-cell and within-cell correlation of cAMPFIRE-L responses to enforced running. n (cells) = 41. Dark gray points indicate median and vertical lines indicate 25^th^ and 75^th^ percentile. Shaded area reflects distribution of data. ***: p < 0.001, dF = 70.2, Welch two-tailed t-test. **e**, Hierarchical clustering of cell-specific responses to 3-min running bouts. **f**, The average (bold lines) and individual (thin lines) traces and quantifications of response amplitudes from cells within each cluster in panel **e**. The percentage of neurons in each category are labeled. From left to right, n (cells) = 7, 8, and 18. Boxes indicate 25th and 75th percentile, middle line indicates median and whiskers indicate range of data excluding outliers. **: p < 0.01 and ***: p < 0.001, dF = 2, one-way ANOVA, Tukey Kramer post-hoc test. **g**, representative images, and the average and individual traces (individual traces corresponding to arrowheads in the image) of running induced response in positively- and negatively-responding cells before and after the administration of yohimbine. **h**, Changes of enforced running-elicited cAMP responses to intraperitoneal injection of the indicated drugs. Yoh.: yohimbine; and prop.: propranolol. From left to right, n (cells) = 6, 3, 4, and 9. Bars indicate mean and error bars indicate s.e.m. Within a group, ***: p < 0.001, paired, two-tailed Student’s t-test; across groups: *: p < 0.05, dF = 7, unpaired, two-tailed Student’s t-test.

To dissect the neuromodulatory pathway underlying the above cell-specific cAMP dynamics, we injected specific neuromodulator receptor antagonists intraperitoneally. We found that the α2 antagonist yohimbine completely abolished the negative responses, whereas the β antagonist propranolol had only moderate, statistically insignificant effect on these responses (Figs. 7g and 7h). These results suggest that the negative responses elicited by enforced locomotion were mediated by the Gi-coupled α2 adrenergic receptors. In contrast, neither yohimbine nor propranolol significantly reduced the positive responses (Fig. 7h), suggesting that another neuromodulator is responsible for the positive cAMP dynamics during animal locomotion. Overall, these results reveal cell-specific, and receptor-specific neuromodulation of L23 pyramidal neurons in the mouse cortex during animal locomotion, which is in part mediated by norepinephrine.

## Discussion

Here, we report cAMPFIREs, a suite of cAMP sensors that are built upon the best-performing available cAMP sensor, and exhibit greatly improved sensitivity, reduced protein aggregation, and corrected subcellular localization. To date, no *in vivo* imaging of cAMP dynamics within individual neurons has been reported. cAMPFIREs allow us to image, for the first time, *in vivo* dynamics of cAMP with cellular and subcellular resolution in mouse cortical neurons during animal behavior. These sensors also function well in *Drosophila* larval neurons (Extended Data Fig. 9), where the growth temperature (25 °C) is substantially different from that in mammals, suggesting that cAMPFIREs will be useful across many biological systems.

Recently, another cAMP sensor, Pink Flamindo has been used to measure cAMP activity in non-neuronal cells in live mice^41^, although the signal remains low and subcellular imaging was not demonstrated. The fact that the optimal maturation temperature of this sensor is below 37 °C^15^ combined with its low response to norepinephrine (Figs. 1d and 1e) may limit its use in neurons in the mammalian brain. In addition, sensors detecting phosphorylation events by PKA have been used for *in vivo* imaging^20^. However, PKA phosphorylation is one step downstream of cAMP and it may also be regulated by phosphatases. Certain cAMP functions are mediated by non-PKA effectors, such as Epac. Direct *in vivo* cAMP imaging, such as that enabled by cAMPFIREs, is needed.

We have developed three cAMPFIREs with different sensitivities that significantly extend the capabilities of previous sensors. In awake mice, we found that cAMPFIRE-M exhibited a significantly elevated baseline in neurons, possibly due to a relatively high tonic level of cAMP. Therefore, the majority of our experiments *in vivo* have employed cAMPFIRE-L. Nevertheless, the sensitivity improvement from the parental sensor to cAMPFIRE-L is critical for *in vivo* applications (e.g., Figs. 6d and 6h). Notably, different cell types may exhibit different levels of cAMP dynamics (e.g., Figure S5). In other cells or other brain regions with lower basal cAMP levels, the higher sensitivity versions of cAMPFIREs may be advantageous.

Our results point to a high tonic level of cAMP *in vivo*, in a large part mediated by norepinephrine. This is consistent with the previous notion that animal wakefulness is associated with increased tones of norepinephrine and acetylcholine^20,37^, and increased activity of PKA^20^. Such tonic levels of cAMP may contribute to the functional and physiological differences between sleep and wakefulness. Interestingly, neuronal somas and dendrites exhibited different tonic levels of basal cAMP activity (Fig. 6d) and different cAMP dynamics to pharmacological manipulations. These results suggest that, between neuronal somas and dendrites, distinct mechanisms, such as differentially expressed adenylyl cyclase subtypes^42^, may be governing the cAMP dynamics, and cAMP-dependent cellular processes may be differentially regulated. In addition, we found cell-specific up or down regulation of cAMP during animal locomotion. This reveals a type of functional heterogeneity in L23 neurons that has not been previously described, possibly mediated by differential expression of neuromodulatory receptor subtypes. Further studies will be required to dissect out the exact function of these different pools of neurons. Nevertheless, given the important roles of cAMP in regulating neuronal excitability and plasticity, this finding may have profound implications in the neuromodulation of brain circuits as well as in animals learning and memory.

In addition to providing information about intracellular signaling mechanisms, cAMP imaging can also be used to infer the functional dynamics of upstream GPCRs and neuromodulators, since three out of four major classes of GPCRs converge onto cAMP. In the mammalian genome, there are ∼1000 GPCRs, which sense tens of different ligands to govern or regulate nearly all aspects of biological processes. Although sensors for detecting several GPCR activators have started to emerge^43^, sensors for the majority of GPCR and their ligands are not available. It is probably also not practical to develop a sensor for each GPCR or for ligands that exist in low-abundance because of the enormous amount of work involved in optimizing each sensor, potential buffering effects, and the theoretically low signal of low-abundance molecules (each molecule can only activate one fluorescent protein). As exemplified in our study, cAMP imaging, together with the appropriate pharmacology or genetic manipulation, may allow one to dissect out the upstream GPCRs and extracellular ligands. This is analogous to using calcium imaging to infer neuronal electric activities. cAMP imaging has the additional advantage that the second messenger signal is vastly amplified from GPCRs, thereby easing detection and reducing the effect of potential buffering. Indeed, we found that expression of cAMPFIREs in neurons of cultured hippocampal slices does not affect neuronal function (Figs. 5e – 5g, and Extended Data Fig. 8), suggesting that the buffering effect to intracellular cAMP, if any, is likely moderate.

Finally, although cAMPFIREs start to enable *in vivo* imaging of neuronal cAMP, they may benefit from further improvements. Further increases in sensitivity are likely unnecessary for intracellular imaging, given that we start to see baseline increases in cAMPFIRE-H even in HEK cells. However, a larger dynamic range may reduce the demand on photon budget, thereby allowing for prolonged imaging or imaging under more challenging conditions, such as in deep brain regions. It may also allow for the *in vivo* visualization of smaller cAMP events or events in smaller neuronal compartments, such as the dendritic spines.

Overall, cAMPFIREs represent a new basis for the application and development of cAMP sensors. Given the importance of cAMP in diverse biological processes and in pharmaceutical development, these sensors will likely have broad applications in both basic and clinical research.

## Methods

All surgical and experimental procedures were performed in accordance with the recommendations in the Guide for the Care and Use of Laboratory Animals, written by the National Research Council (US) Institute for Laboratory Animal Research, and were approved by the Institutional Animal Care and Use Committee (IACUC) of the Oregon Health and Science University (#IP00002274).

### Plasmid Constructs

Constructs were made using standard mutagenesis and subcloning methods, or by gene synthesis (Genewiz). All previously unpublished constructs and their sequences will be deposited to Addgene.

### Cell culture and transfection

HEK-293 cells (ATCC # CRL-1573) were maintained in 100-mm cell culture dishes (Fisher Scientific, #FB012924) at 37°C with 5% CO_2_ in minimal essential medium (ThermoFisher #11095-080) with the addition of 10% fetal bovine serum and penicillin-streptomycin. All cells from ATCC have been authenticated by morphology, karyotyping, and PCR based approaches. These include an assay to detect species-specific variants of the cytochrome C oxidase I gene (COI analysis) to rule out inter-species contamination, and short tandem repeat (STR) profiling to distinguish between individual human cell lines to rule out intra-species contamination. These cells are also tested for mycoplasma by ATCC. Cell aliquots were kept frozen in liquid nitrogen until use, and were further authenticated based on their morphology. Once thawed, each aliquot of cells will be passed and used for no more than 4 months. To image transfected HEK cells under our two-photon microscope, cells were subcultured onto glass coverslips (hand-cut to ∼5 x 5 mm) coated with 0.1 mg/mL Poly-D-Lysine (Millipore-Sigma, #27964-99-4) in 35-mm cell culture dishes (Corning, #CLS430165).

For sensor expression, constructs in mammalian expression plasmids (0.5 – 3 µg/35-mm dish) were transiently transfected using Lipofectamine-2000 (ThermoFisher, #11668030) according to the manufacturer’s instructions with the exception that only half of the recommended amount of the reagent (5 µl) was used. Imaging was performed at two days post-transfection in a chamber perfused with gassed (95% O_2_/5% CO_2_) artificial cerebral spinal fluid (aCSF) containing (in mM) 127 NaCl, 25 NaHCO_3_, 25 D-glucose, 2.5 KCl, 1.25 NaH_2_PO_4_, 2 CaCl_2_, and 1 MgCl_2_.

### Immunoprecipitation

HEK-293 cell culture and construct transfection were performed as above, with the exception that transfections were performed in 100-mm cell culture dishes to increase total protein yield. Two days post-transfection, cells were lysed with Triton-Immuno Precipitation Buffer (IPB) containing 1X PBS supplemented with 3 mM EDTA, 0.1 mM PMSF, 1X Aprotinin, 2% BSA, and 1% Triton X100, and spun down at > 100,000g for 10 minutes. The supernatant was combined with protein-A agarose beads (ThermoFisher #20333) and a mouse monoclonal anti-myc antibody, and rotated at 4°C overnight. Beads were then spun down, and washed 2x with Triton-IPB and then 2x with IPB (Triton-IPB without the addition of Triton X100), with a 3-5 minute rotation and spin-down between each wash. After the last wash, beads were resuspended in IPB and stored at 4°C. Notably, we have empirically found that the addition of 1-2% BSA is critical to maintain the dynamic range of the precipitated sensors, possibly by reducing protein denaturing in low protein environments. To quantify sensor response, beads were loaded in a chamber perfused with gassed (95% O_2_/5% CO_2_) aCSF with 25 mM HEPES, 1 mM ascorbic acid, and 2 mM (final concentration) Mg^2+^. To hold each bead under perfusion, negative pressure was applied to suction each bead to a borosilicate glass pipette (tip diameter ∼20-60 μm).

### Hippocampal slice culture and transfection

Hippocampi were dissected from P6 – P7 rat pups of both sexes. Sections (400 µm) were prepared using a chopper in dissection medium containing (in mM) 1 CaCl_2_, 5 MgCl_2_, 10 glucose, 4 KCl, 26 NaHCO_3_, and 248 sucrose, with the addition of 0.00025% phenol red. The slices were then seeded onto a cell culture insert (Millipore, #PICM0RG50) and cultured at 35°C with 5% CO_2_ in 7.4 g/L MEM (ThermoFisher, #11700-077) with the addition of (in mM unless labeled otherwise): 16.2 NaCl, 2.5 L-Glutamax, 0.58 CaCl_2_, 2 MgSO_4_, 12.9 D-glucose, 5.2 NaHCO_3_, 30 HEPES, 0.075% ascorbic acid, 1 mg/mL insulin, and 20% heat-inactivated horse serum. Slice media was refreshed every 2-3 days after seeding replacing ∼60% of the culture media.

Transfection of cultured slices was accomplished using the biolistic method (Bio-Rad Helios gene gun) 10 to 20 days after seeding. In short, slices were bombarded with biolistic particles created by coating 1.6 μm gold particles (Bio-Rad, #165-2262; ∼1 μg DNA/mg gold) with constructs in mammalian expression plasmids. Cultured slices were imaged at 2-3 days post-transfection in a chamber perfused with gassed (95% O_2_ / 5% CO_2_) aCSF that contained 4 mM CaCl_2_ and 4 mM MgCl_2_.

### *In utero* electroporation

C57BL/6 mice (Charles River, or home-bred within 1 generation from Charles River breeders) were used for acute slices and *in vivo* experiments. To express sensors in mice, *in utero* electroporation was performed at E15.5 by injecting plasmid (1 μL/embryo, concentration ∼2-3 μg/μL, mixed with a 0.2% final concentration of fast green for visualization) into the lateral ventricle of mouse embryos, which were then electroporated with five 100-ms pulses (35V, 1 Hz) using an electroporator (BEX #CUY21). To achieve expression in the somatosensory cortex, the positive terminal of the electroporator was positioned above the right parietal field before the pulses were delivered.

### Patch-clamp electrophysiology

Whole-cell patch-clamp was performed in CA1 pyramidal neurons from organotypic cultured slices. Sequential recordings were done in transfected neurons and in adjacent untransfected neurons (control) within 50 µm in the same slice. Voltage-clamp recordings were performed using a MultiClamp 700B amplifier (Molecular Devices) controlled with custom software written in MATLAB. Electrophysiological signals were filtered at 2 kHz before being digitized at 20 kHz. Slices were perfused at room temperature with gassed aCSF containing 4mM CaCl_2_ and 4 mM MgCl_2_. Recording pipettes (3–5 MΩ), were pulled from borosilicate glass (G150F-3; Warner Instruments) using a model P-1000 puller (Sutter Instruments). Series resistance was 10–25 MΩ. The internal solution contained (in mM): 126 Cs-gluconate, 10 CsCl, 10 HEPES, 5 Na-phosphocreatine, 0.5 Na-GTP, 4 Mg-ATP, 10 TEA-Cl, 5 EGTA and 4 QX-314 bromide with an osmolarity of 295-300 mOsmol/kg and pH ∼7.2 adjusted with CsOH. The junction potential was calculated to be −14 mV, as calculated using JCal from Clampex software (Molecular Devices). Voltages were not corrected for the theoretical liquid junction potentials.

AMPA/NMDA responses were evoked by extracellular stimulation of the Schaffer collaterals using a theta-glass (20-50 µm tip) filled with aCSF and positioned at 150 µm lateral from somas. Short pulses (0.1 ms) were applied using an A365 stimulus isolator (World Precision Instruments) to evoke monosynaptic responses. GABA_A_ synaptic currents were blocked by adding 100 µM picrotoxin in the bath. To prevent recurrent activities, connections between CA1 and CA3/subiculum were cut and 4 µM 2-chloroadenosine was added to reduce polysynaptic responses. In voltage-clamp mode, cells were held first at a holding potential (V_H_) of +55 mV to record NMDA component and at V_H_ = −60 mV to record AMPA component. 10 trials were recorded using an interstimulus interval of 10 s. AMPA/NMDA ratio were determined as the ratio between peak amplitude of the response evoked at V_H_ = −60 mV over NMDA component measured at +55 mV as the mean current at 140-160 ms after stimulation. Paired-pulse ratios were calculated as the ratio between peaks (2^nd^ peak/1^st^ peak) from traces containing two evoked responses separated by 70 ms at V_H_ = −70 mV in voltage-clamp mode. 10 trials were recorded using an interstimulus interval of 10 s. For mEPSCs recordings, 1 μM TTX, 10 μM CPP, and 100 μM Picrotoxin were added to the bath to block action potentials, NMDA and GABA_A_ receptors. 5-minute traces were recorded and mEPSCs events were detected using a template matching built-in feature in Clampfit (Molecular devices).

### Acute slice preparation

Mice with neurons transfected via *in utero* electroporation were transcardially perfused with ice-cold, gassed aCSF containing (in mM) 127 NaCl, 25 NaHCO_3_, 25 D-glucose, 2.5 KCl, 1.25 NaH_2_PO_4_, 2 CaCl_2_, and 1 MgCl_2_. The brain was then resected and coronally sliced using a vibratome (Leica VT1200s) in an ice-cold, gassed choline-cutting solution containing (in mM): 110 choline chloride, 25 NaHCO_3_, 25 D-glucose, 2.5 KCl, 7 MgCl_2_, 0.5 CaCl_2_, 1.25 NaH_2_PO_4_, 11.5 sodium ascorbate, and 3 sodium pyruvate. The slices were then incubated in gassed aCSF at 35°C for 30 minutes and subsequently kept at room temperature for up to 8 hours.

### Cranial window surgeries

For *in vivo* imaging, a glass window was installed on the skull of the mouse between 30 – 60 days postnatal. The open skull surgery was performed under inhaled isoflurane anesthesia (4% to induce, ∼1.5-2% to maintain). Briefly, the skull was exposed and cleaned using a Q-tip and all connective tissue was cleared. Next, a circular ∼3.5 mm craniotomy was made, using an air-powered high-speed dental handpiece (Maxima Pro 2)and a ¼ inch carbide burr (Henry Schein #5701072), above the somatosensory cortex. The window was implanted into the craniotomy site and adhered to the skull with dental acrylic. Last, a steel plate was adhered to the skull (for head fixation) using dental acrylic. Mice were allowed to recover for two weeks prior to imaging and were treated with pre-operative dexamethasone (4 mg/mL) and post-operative carprofen (0.5 mg/mL) to reduce inflammation.

### Two-photon and 2pFLIM imaging

The *in vitro* two-photon microscope was built as previously described^44^ and the *in vivo* two-photon microscope was built based on the open-access design of the Modular In vivo Multiphoton Microscopy System (MIMMS) from Howard Hughes Medical Institute Janelia Research Campus (https://www.janelia.org/open-science/mimms). Both two photon microscopes were controlled by the ScanImage software^44^ Vidrio). Fluorophores were excited with a pulsed 80 MHz Titanium-Sapphire laser at the following wavelengths: (850 nm for Epac1-camps, Epac2-camps300, ICUE3, Epac-S^H187^, and all cAMPFIRE variants; 960 nm for Flamindo2 and cAMPr; and 990 nm for Pink Flamindo and R-FlincA). The fluorescence emission for these were unmixed using a dichroic mirror and band-pass filters. Specifically, Semrock FF511-Di01, Semrock FF01-483/32 and Semrock FF01-550/49 were used for Epac1-camps, Epac2-camps300, ICUE3, Epac-S^H187^, and all cAMPFIRE variants. The Chroma 565DCXR dichroic was used for Flamindo2, Pink Flamindo, cAMPr, and R-FlincA, with a Semrock FF01-630/92 barrier filter used for Pink Flamindo and R-FlincA, and a Chroma HQ510/70 barrier filter for cAMPr and Flamindo2.

2pFLIM was carried out as recently described^20,36^. Briefly, four additional hardware components were integrated into the two-photon setup. A photodiode (Thorlabs FDS010) was used to detect the arrival of the laser pulses, and a fast, cooled photomultiplier tube (Hamamatsu H7422PA-40 or H10769PA-40) was used to detect the arrival of the donor-emitted photons. These arrival times were compared using a TCSPC-730 or SPC-150 (Becker and Hickl) time-correlated single photon counting board. Where needed, a frequency to analog converter (Becker and Hickl HPM-CON-02) was used to split the signals into two outputs: one with an unaltered waveform for lifetime measurements and another with a slower output (∼2 µs) for conventional two-photon imaging. Data acquisitions were controlled by a modified version of FLIMimage, which was written in MATLAB and was kindly provided by Dr. Ryohei Yasuda. FLIMimage functions as an integrated add-on to ScanImage.

For *in vivo* imaging experiments, animals were head-fixed and placed on a one-dimensional treadmill which, except during locomotion-related experiments, was fixed. Isoflurane (1.5%) was delivered via direct inhalation through a nose cone. All other pharmacological reagents were administered via intraperitoneal (IP) injection.

### Image analysis

Data analyses were performed using custom software written in MATLAB or Python. In particular for 2pFLIM analyses, we used a software suite called FLIMview written in MATLAB^20^. Where appropriate, regions of interest were drawn to isolate HEK cell, somatic or dendritic signals from contamination by background photons or adjacent cells/structures. For ratiometric imaging, individual channels were background subtracted based on a nearby background ROI before calculating the ratio. Fluorescence lifetime was approximated by mean photon emission time τ:

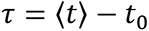

such that t reflects the emission time of individual photons in the measurement window and t_0_ reflects the timing of laser pulses. t_0_ is a fixed property of a given hardware configuration and is measured separately under ideal imaging conditions. Because the measurement window (t_w_) is finite (≤ 10 ns passing t_0_ due to laser pulse repetition and single photon counting hardware properties), the measured τ (τ_apparent_) is slightly smaller than the real value. For example, for a τ of 2.0 ns and a t_w_ of 9.0 ns, the τ_apparent_ is 1.90 following the equation below:

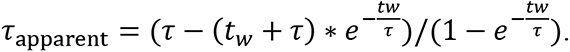

### Analysis of *in vivo* data

Quantification of fluorescence lifetime under control and pharmacologically manipulated conditions was conducted by averaging ≥ 3 representative time points. For Fig. 7d, Pearson’s correlation coefficient was used to represent the correlation of the time-dependent response to enforced running during the three-minute running window.

Correlation was calculated between repeated trials from a single cell or from average response across trials between different cells. For Fig. 7e, hierarchical clustering was performed on the average of responsive bouts for each cell. Euclidean distance and the Ward linkage metric were used after careful examination of the efficacy of multiple linkage metrics and clustering algorithms. Output clusters were similar across both hierarchical and non-hierarchical (K-mean) clustering algorithms and the hierarchical method was chosen so as to reduce bias from predetermining cluster number. Prior to clustering, cells with data from five or fewer running bouts were excluded (six cells) as was a single cell with no response to running. A bout was deemed responsive if the integral of the peak response surpassed the integral of the standard deviation of the baseline data in the same time domain. The averaged baseline standard deviation across all cells was 0.021 ns. Graphs showing cell lifetime over time were smoothed using a three point moving average.

### Generation of fly transgenic lines

cDNAs for cAMPFIRE-L, cAMPFIRE-M, cAMPFIRE-H, and mTurquoise2 were amplified from mammalian plasmids by PCR and cloned into the pUASTattB plasmid via In-Fusion® HD cloning (Takara Bio USA). td-cpVenus was amplified by fusion PCR and cloned into the pUASTattB plasmid at the HindIII and XbaI restriction sites. The resulting constructs were individually microinjected (250 ng/µL, Eppendorf FemtoJet) into embryos of yv nos-phiC31; P{CaryP}attP40; (BL# 25709) flies to generate stable transgenic lines.

### Live Imaging of flies

Embryos were collected for 6-7 hr at 25 °C on molasses-based cornmeal food and grown at 25 °C until late 3rd instar. Larvae were then dissected and imaged in hemolymph-like 3 (HL3) saline with 10 mM glutamate to suppress muscle movement (in mM: 70 NaCl, 5 KCl, 1.5 CaCl_2_, 20 MgCl_2_, 5 trehalose, 115 sucrose, 5 HEPES, 10 glutamate). To expose the central nervous system (CNS) for imaging, larvae were immobilized by insect pins, cut along the dorsal midline, and splayed open to expose internal tissues. Internal tissues were carefully removed from the anterior half of the larvae, taking care not to touch the CNS and keeping the larvae as intact as possible. Live imaging was performed using a Leica SP5 confocal system equipped with an acousto-optical beam splitter and an HC Fluotar L 25X/0.95 W VISR immersion objective (Leica). Fluorescence was excited by a 405 nm laser and collected by two photomultiplier tubes set to 440-500nm (mTurquoise2) and 575-700 nm (td-cpVenus signal). The pinhole was set to 3 AU to maximize collection of emitted photons. Image stacks were acquired every 10 seconds at a scan speed of 1400 Hz with 2X line averaging using LAS AF 2.7.3 (Leica).

### Analysis of fly imaging data

All images were analyzed in ImageJ (NIH). Briefly, drift correction was performed using the Correct 3D Drift plugin with multi time scale computation. Background for each channel was estimated as the mean fluorescence during the first 20 time points within an ROI drawn over a background region of the image. Background was then directly subtracted for each channel before calculating a maximum intensity projection. An ROI containing the cell body was defined based on the first image using Intermodes Thresholding followed by Binary Erosion. The mean fluorescence intensity of the ROI was calculated for each timepoint and recorded in Apple Numbers. The FRET ratio was then calculated as mTurquoise/td-cpVenus. Graphs showing FRET ratios over time were smoothed using a 5-point moving average and normalized to the initial FRET value.

### Data presentation and statistical analysis

Quantification and statistical tests were performed using custom software written in MATLAB. Averaged data are presented as mean ± s.e.m., unless noted otherwise. In box plots, lower and upper box lines indicate 25th and 75th percentile, respectively, with middle line indicating median and whiskers indicating total range of data, not including outliers. All measurements were taken from different cells. “n” refers to the number of cells unless noted. In HEK cell experiments, most (>90%) experiments measured two cells simultaneously per coverslip, with the remaining measuring either three or four cells. In cultured and acute slice experiments, all neurons came from different slices. All experiments were repeated for at least two, typically more, independent transfections (*in vitro*) or mice (*in vivo*). Unless otherwise noted, the Bartlett test was used to determine if variances were equal, and the Shapiro Wilk test was used to determine if data were normally distributed. When the assumptions of normality and homoscedasticity were met, a two-tailed Student’s t-test or one-way ANOVA followed by a Tukey-Kramer post-hoc test were used. When the assumption of homoscedasticity was not met, a Welch’s t-test was used. In all figures, *: p < 0.05, **: p < 0.01, and ***: p < 0.001.

## Acknowledgements

We thank Dr. Ryohei Yasuda (Max Planck Florida) for 2pFLIM acquisition software; Dr. Jin Zhang (University of California, San Diego) for ICUE3 cDNA; Dr. Kees Jalink (Netherlands Cancer Institute) for Epac-S^H187^ cDNA; Drs. Laurinda Jaffe (University of Connecticut) and Viacheslav Nikolaev (University of Hamburg) for Epac1-camps and Epac2-camps300 cDNA; Drs. Lei Ma and Vivek Unni and Ms. Sydney Boutros at Oregon Health & Science University and Josh Melander at Stanford University for training and advice on *in vivo* surgery; Dr. Kevin Wright (Vollum Institute) for immunoprecipitation reagents; Drs. John Williams, Arpiar Saunders, Bart Jongbloets, Michael Muniak, and Lei Ma, Ms. Yang Chen, and the entire Zhong lab and Mao lab at Oregon Health and Science University for helpful comments and discussions. Fly stocks from the Bloomington Drosophila Stock Center (NIH P40OD018537) were used. This work was supported by two NIH BRAIN Initiative awards R01NS104944 and RF1MH120119 (H.Z. and T.M.), and an NINDS R01 grant R01NS081071 (T.M.). The study on *Drosophila* was supported by an NIH R01 grant R01NS104299 (B.Y.).

## Author Contributions

HZ conceived the project. CIM, TM, and HN designed the experiments. MW, CIM, and HZ performed structural analyses of Epac. CIM performed *in vitro* experiments with assistance from HN and LBE, except that CCC performed electrophysiological recordings. CIM and LBE carried out mouse *in vivo* imaging experiments. MZ prepared and maintained cultured slices and mouse husbandry, and performed IUE surgeries. EC and BY performed all experiments and data analysis associated with fly imaging, and EC and BY wrote the corresponding legends and methods. CIM, LBE, TM, and HZ analyzed the data and wrote the manuscript, with edit and comments by all authors. TM and HZ secured the funding and supervised the project.

## Data availability

All previously unpublished sensor constructs and their corresponding sequences will be deposited to Addgene (http://addgene.org). Source data will be provided with this paper.

## Code availability

Custom MATLAB codes will be made available upon request.

## Extended Data Figures and Table

**Extended Data Figure 1.**
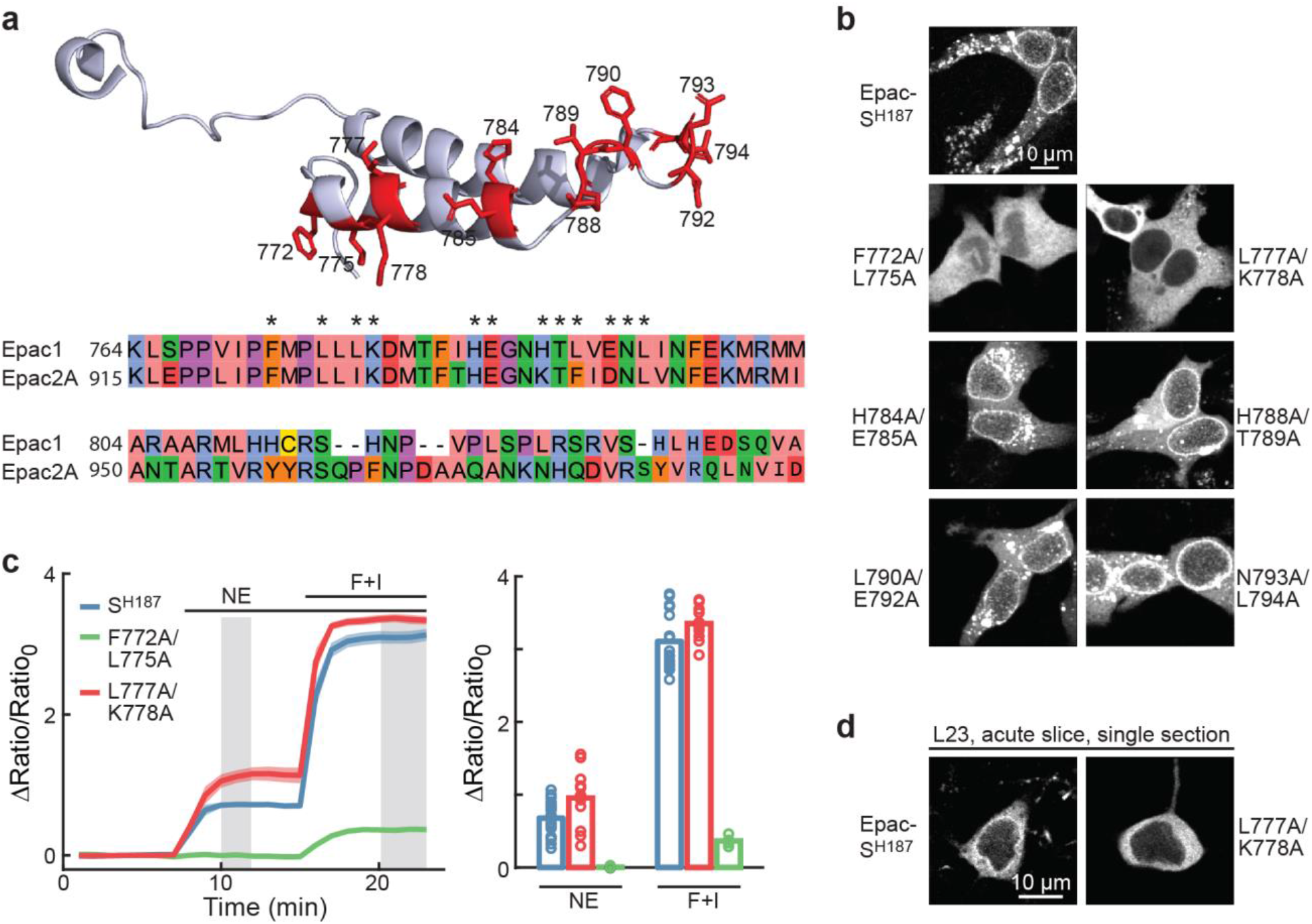
Correcting the perinuclear localization of the Epac-S^H187^ sensor. **a**, Crystal structure of the NPL sequence in Epac2A (upper) and the alignment of NPL sequences between Epac1 with Epac2. The positions tested are highlighted in the structure (red) and at the alignment (asterisks). **b**, Representative images of the indicated mutants compared to Epac-S^H187^ in HEK cells. **c**, Response time course (left) and quantification of the response of indicated constructs to the indicated stimuli in HEK cells. From left to right, n = 20, 13, and 11. Dark lines and bars indicate mean. Shaded area indicates s.e.m. **d**, Representative images of the indicated mutants compared to Epac-S^H187^ in L23 pyramidal neurons. Single two-photon optical sections are shown.

**Extended Data Figure 2.**
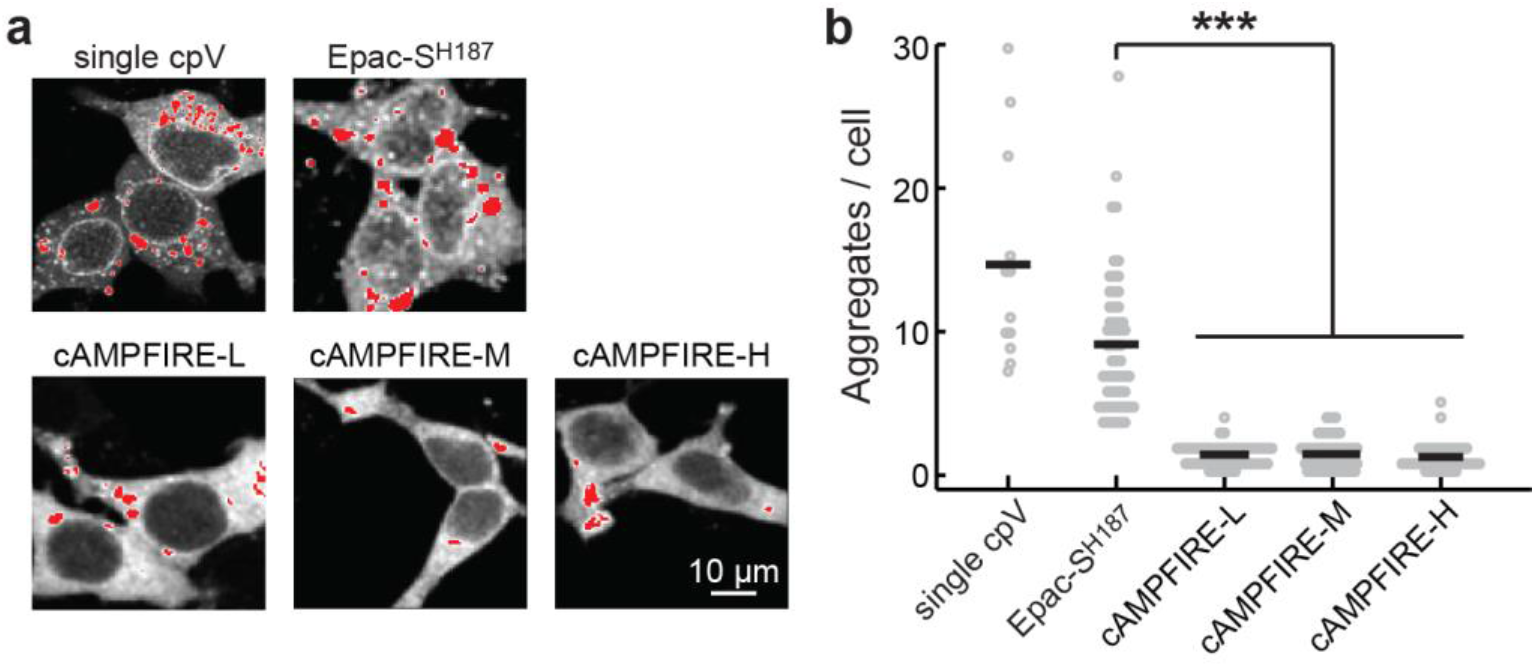
Localization-correcting mutations reduce the tendency of the sensor to aggregate. **a** & **b**, Representative images (**a**) and quantifications (**b**) of the number of aggregates/cell resulting from the indicated constructs. To ease the visualization of aggregates, saturated pixels (for display only) are showed in red. Single cpV: a sensor variant that is the same as Epac-S^H187^ with the exception that it only has a single cpVenus as the FRET acceptor. From left to right, n (cells) = 12, 58, 53, 48, and 41. Lines indicate mean. ***: p < 0.001 for all comparisons, two-tailed Welch test.

**Extended Data Figure 3.**
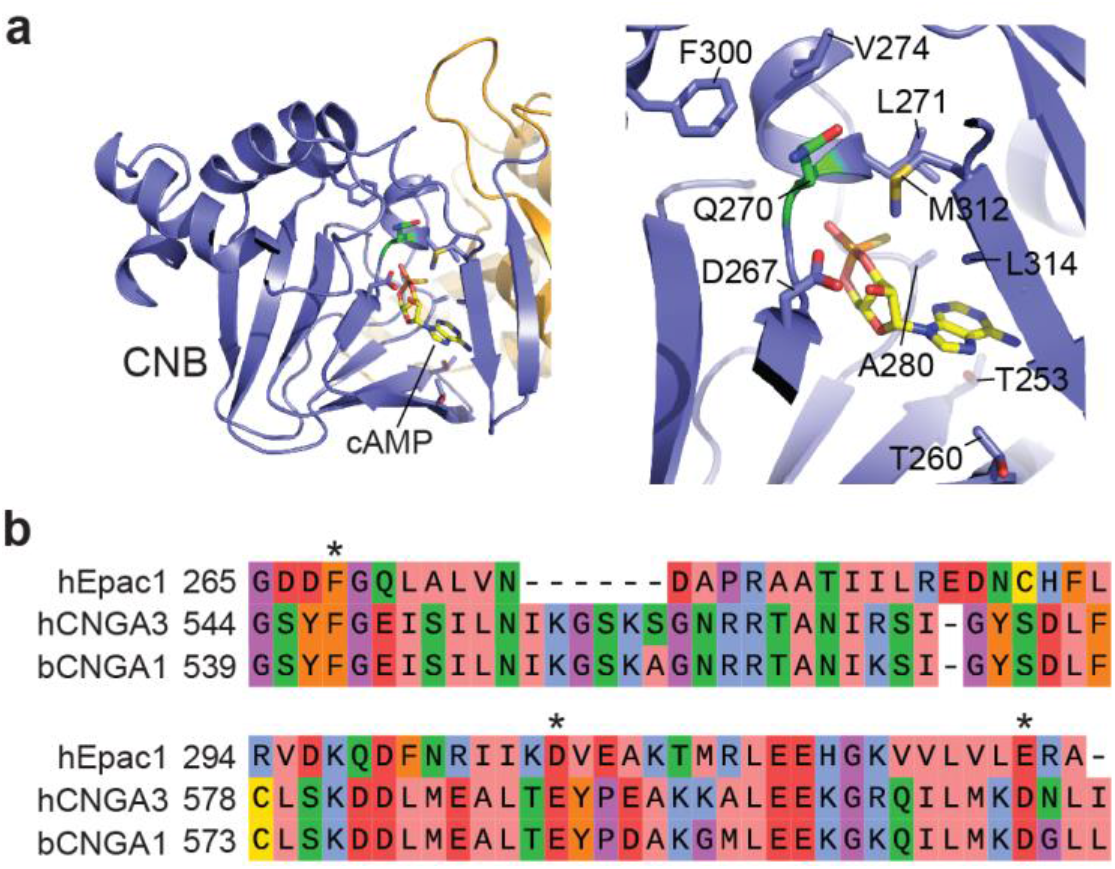
Candidate residues for tuning the affinity of cAMP binding. **a**, Crystal structure analyses of the cAMP binding site of Epac2A suggests 10 positions that potentially affect the cAMP binding affinity. **b**, Alignment of a portion of the CNB domain of Epac1 with those of the CNG channel CNGA2 and CNGA3. Known positions at which mutations affect the affinity of the CNG channels to their ligand are shown (asterisks).

**Extended Data Figure 4.**
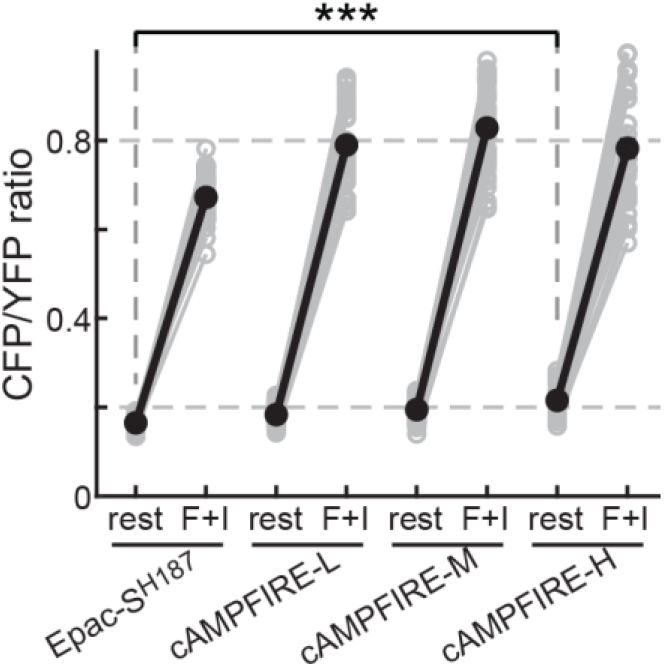
Responses of cAMPFIREs in absolute CFP/YFP ratios. The resting CFP/YFP ratios were: 0.165 ± 0.001 for Epac-S^H187^, and 0.221 ± 0.003 for cAMPFIRE-H; p<0.001. From left to right, n (cells) = 60, 51, 51, and 55. Black points indicate mean. ***: p < 0.001, two-tailed Welch test.

**Extended Data Figure 5.**
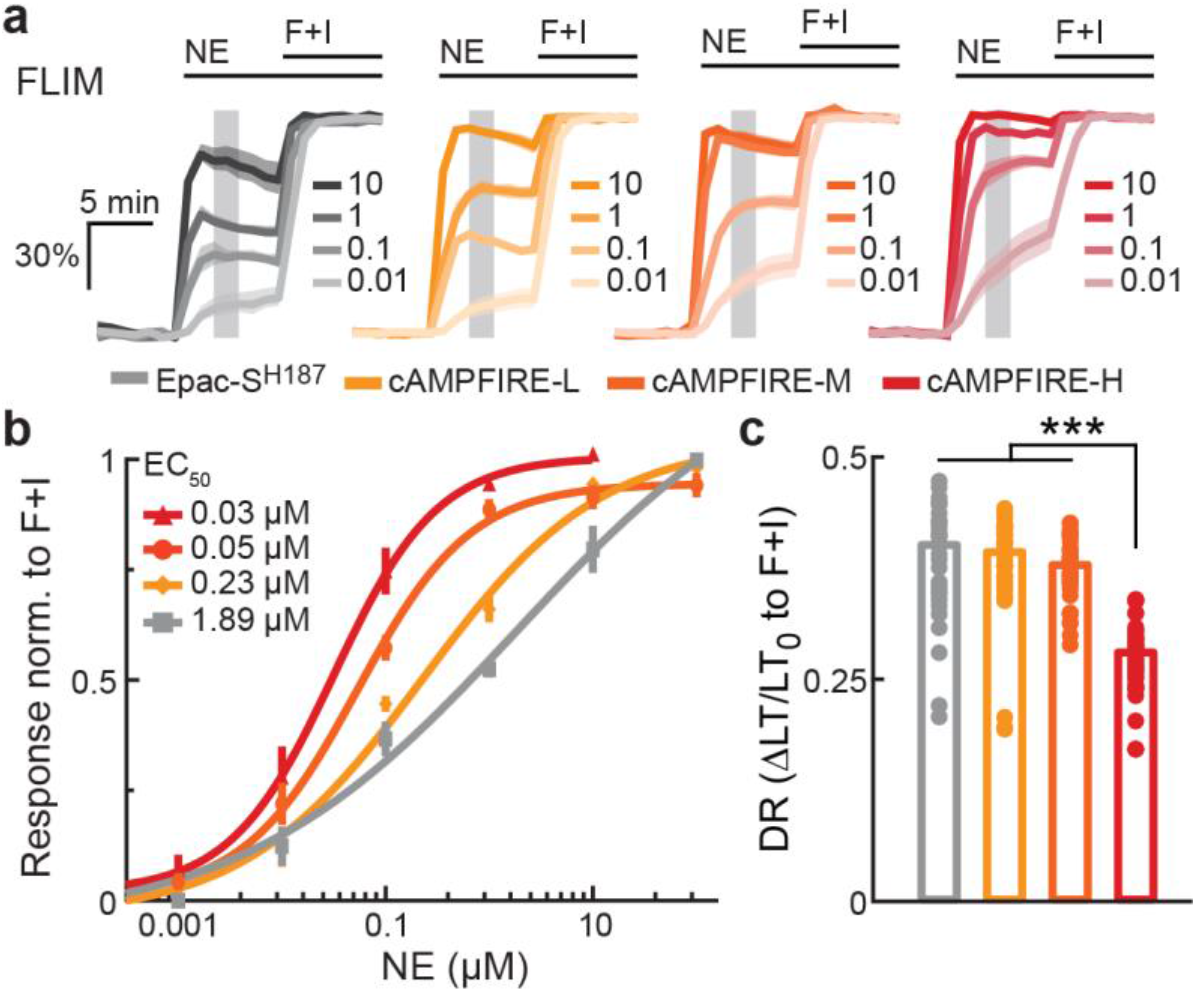
2pFLIM characterizations of cAMPFIREs in HEK cells. **a** & **b**, 2pFLIM response traces (**a**), and NE dose-response curves fitted by the Hill equation (**b**), of Epac-S^H187^ and cAMPFIRE-L, M, and H in HEK cells. n = 4 – 14 for each sensor and concentration data point. Dark lines and shaded area in panel **a,** and points and error bars in panel **b** indicate mean and s.e.m., respectively. **c**, 2pFLIM response amplitudes of each sensor elicited by 25 µM forskolin and 50 µM IBMX. From left to right, n (cells) = 40, 51, 49, and 49. Bars indicate mean. ***: p < 0.001 for all comparisons, two-tailed Welch test.

**Extended Data Figure 6.**
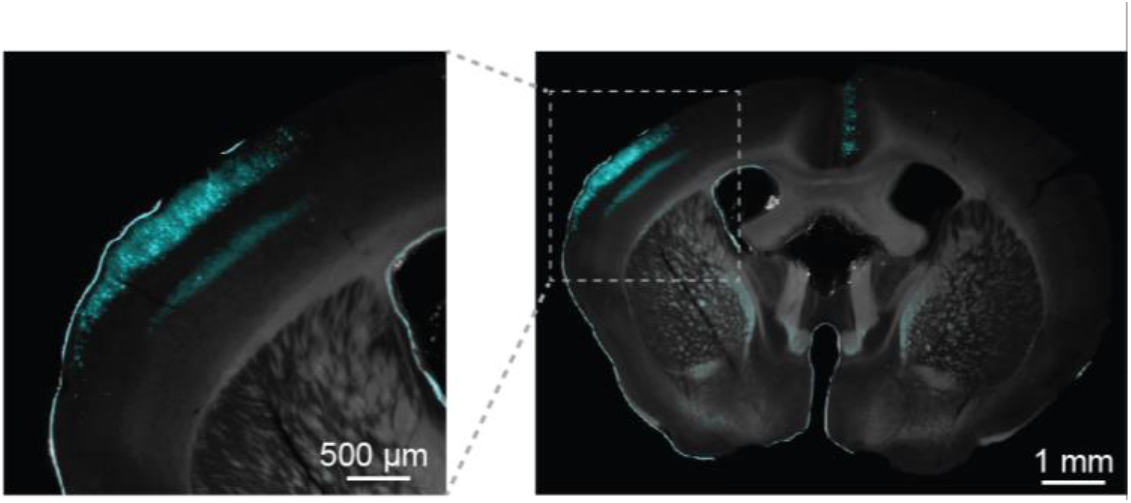
*In utero* electroporation transfection of L23 neurons in the somatosensory cortex. Example brain slice images from a mouse transfected with cAMPFIRE-L via *in utero* electroporation. The sensor-transfected neurons are shown in cyan.

**Extended Data Figure 7.**
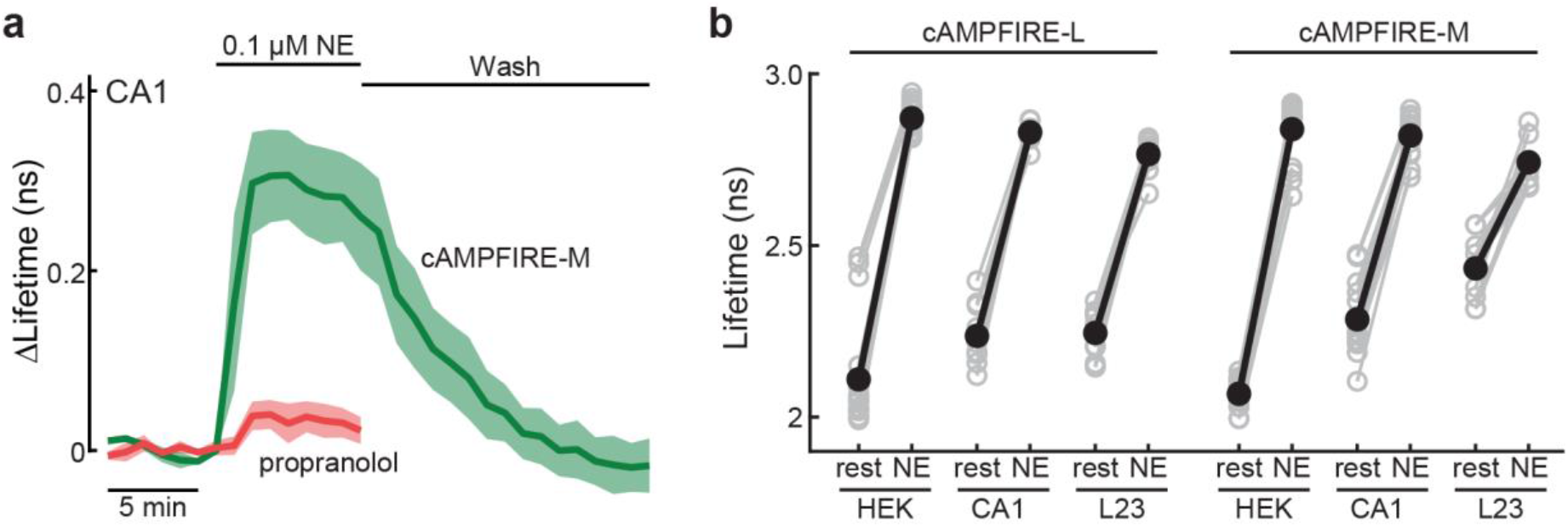
Characterizations of cAMPFIRE responses in neurons. **a**, Response traces of cAMPFIRE-M in CA1 neurons in cultured hippocampal slices to 0.1 µM norepinephrine in the absence or presence of 1 µM propranolol, followed by wash. n (cells) = 4 for both. Dark lines indicate mean and shaded area indicates s.e.m. **b**, 2pFLIM responses in absolute values of cAMPFIRE-L and cAMPFIRE-M to 1 µM norepinephrine in different cell types, indicating that different cell types exhibit different baseline and dynamic ranges. From left to right, n (cells) = 21, 10, 12, 18, 16, and 10. Black points indicate mean.

**Extended Data Figure 8.**
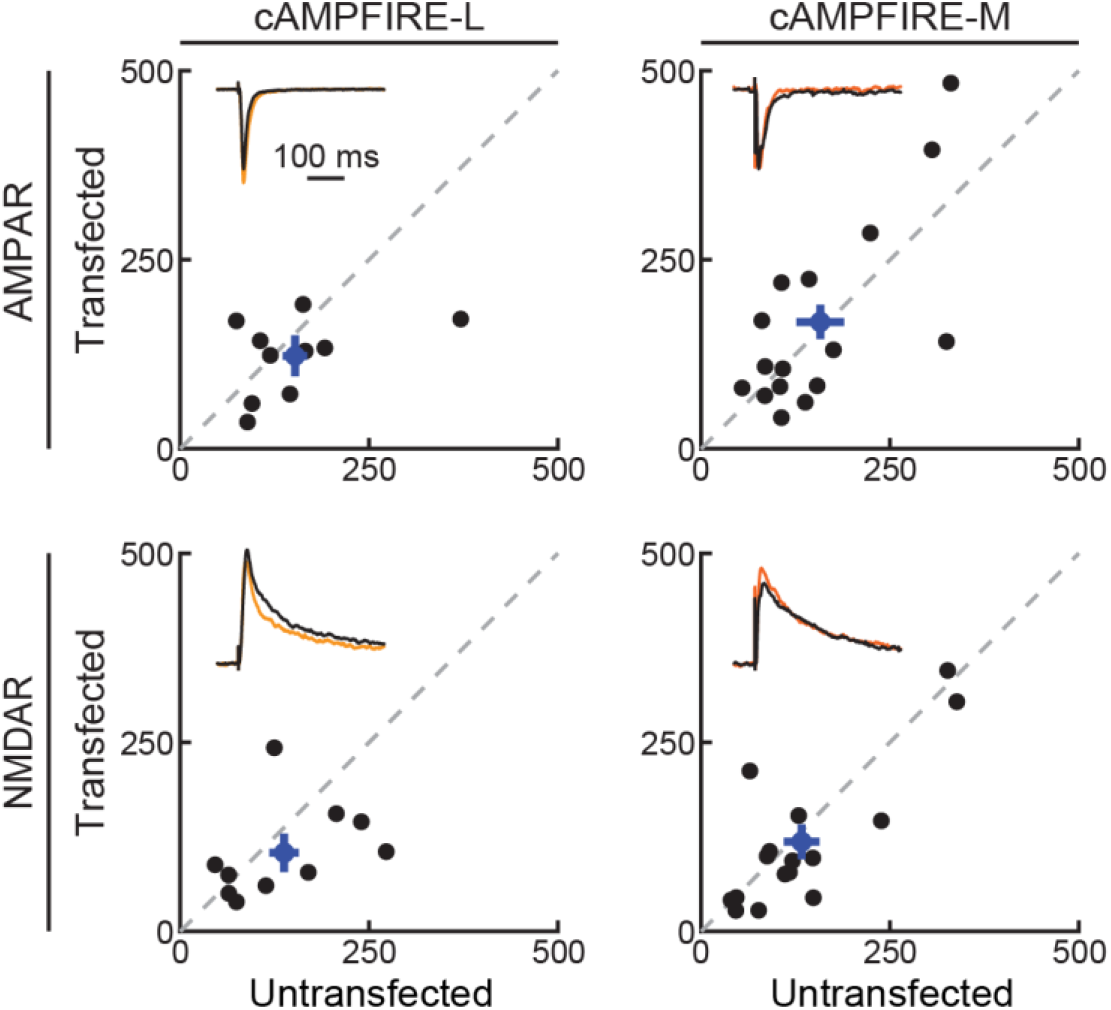
Additional electrophysiological characterizations of cAMPFIREs in CA1 neurons. Quantification of the elicited AMPAR and NMDAR currents (in pA) of CA1 neurons transfected with cAMPFIRE-L or cAMPFIRE-M compared to adjacent untransfected control neurons. For both, n = 10 neuronal pairs for cAMPFIRE-L and 16 for cAMPFIRE-M. Blue points and error bars indicate mean and s.e.m., respectively.

**Extended Data Figure 9.**
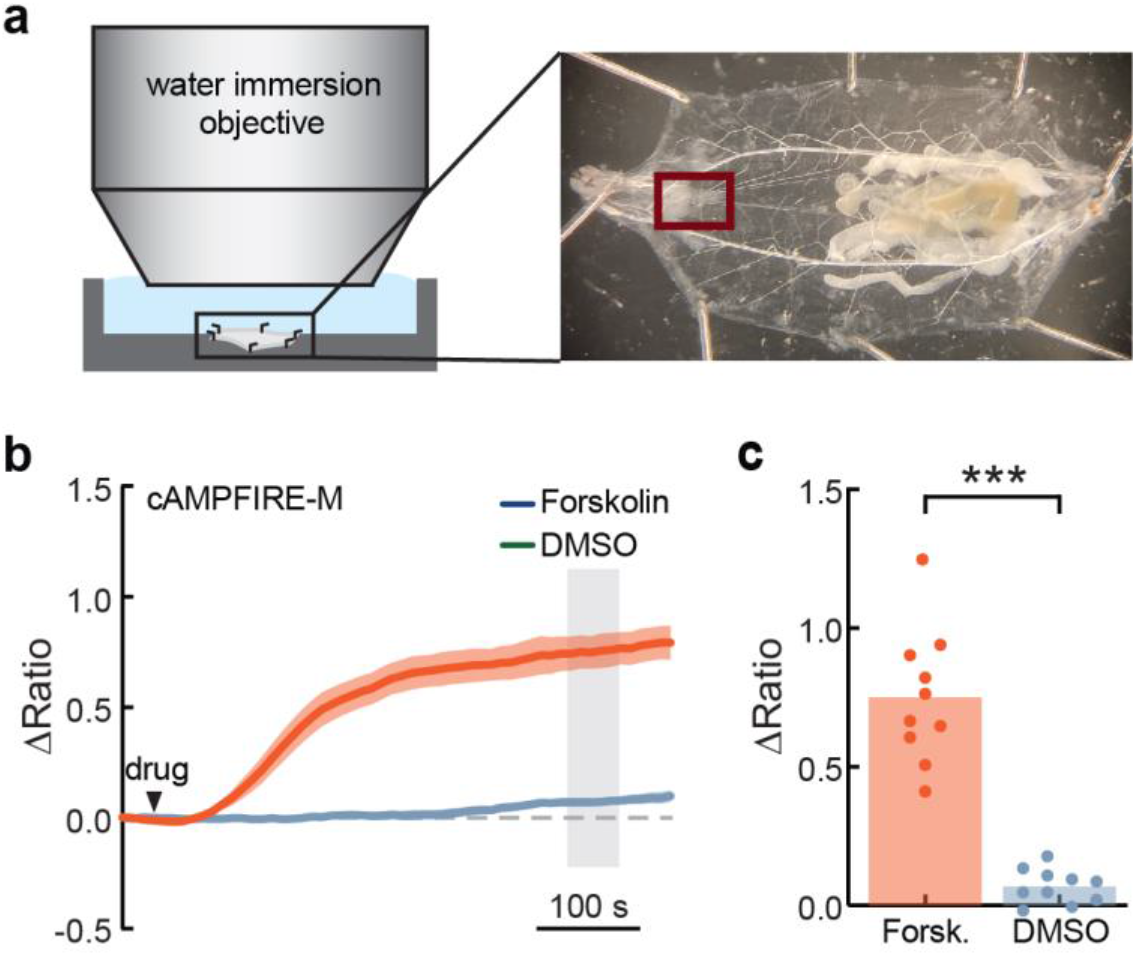
cAMPFIRE-M detects cAMP responses in *Drosophila* larvae. **a**, Schematic showing experimental setup. Larvae were dissected to expose the intact central nervous system (boxed region) and imaged with a water immersion objective. **b**, Responses of abdominal leucokinin neuronal somas expressing cAMPFIRE-M following application of forskolin (40 µM) or DMSO-only control. Dark lines indicate mean, shaded area indicates s.e.m. Arrowhead indicates time of bath application. **c**, Maximum change in FRET ratio for forskolin-treated or DMSO-treated larvae, calculated as an average value from the time window shaded in panel **b**. Bars indicate mean. ***: p < 0.001, Wilcoxon Rank-Sum test. n = 10 larvae per condition.

**Extended Data Table 1:** *Drosophila* lines used in this study.

| ***Drosophila* line** | **Source** | **Identifier** |
| --- | --- | --- |
| w; LK-GAL4 | Bloomington | 51993 |
| yv nos-phiC31; P{CaryP} attP40; | Bloomington | 25709 |
| w; UAS-CAMPFIRE-H [attP40]; | This study | N/A |
| w; UAS-CAMPFIRE-M [attP40]; | This study | N/A |
| w; UAS-CAMPFIRE-L [attP40]; | This study | N/A |
| w; UAS-cp-tdVenus [attP40]; | This study | N/A |
| w; UAS-mTurquoise2 [attP40]; | This study | N/A |

